# Endothelin Signaling via EDNRB receptor Reduces Proliferation and Promotes Proneural-to-Mesenchymal Transition in Gliomas

**DOI:** 10.1101/2025.09.05.674570

**Authors:** D Pineau, L Garcia, H Arnold, A Hanžek, M Arrieta, L R Gauthier, C Granotier-Beckers, F D Boussin, A Herbet, M Hautière, V Asei-Ceschino, C Jacques, L Brard, T Harnois, V Coronas, B Constantin, A Chatelier, J Chemin, S Urbach, M Seveno, S Hideg, C Ripoll, K Aguilar-Cazarez, M Zheng, Guo-Hao Huang, Sheng-Qing Lv, L Zhang, P Rondard, L Prezeau, JP Pin, H Duffau, L Bauchet, V Rigau, F Denat, C Truillet, D Boquet, JP Hugnot

**Author notes:** **Correspondance:** JP HUGNOT, IGF-Institut de Génomique Fonctionnelle, INSERM U 1191 - CNRS UMR 5203-Montpellier University, 141, rue de la Cardonille, 34094 Montpellier Cedex 5 France.

## Abstract

Diffuse gliomas are incurable primary brain tumors encompassing three histo-molecular subtypes: glioblastomas (GB), astrocytomas, and oligodendrogliomas. The latter two harbor IDH1 mutations and exhibit slower progression than glioblastomas. Diffuse gliomas are composed of highly plastic tumor cells capable of transitioning between astrocyte-like, oligodendrocyte-like, progenitor-like, and mesenchymal-like states, driven by genetic alterations and microenvironmental cues. The proneural-to-mesenchymal transition (PMT), associated with increased malignancy, is notably influenced by cytokines in the tumor microenvironment. Endothelin cytokines (ET-1, ET-2, ET-3), primarily secreted by vascular cells, regulate not only vascular tone but also astrocyte and neural stem cell proliferation via the G-protein-coupled receptors EDNRA and EDNRB. Prior studies using serum-cultured glioma lines suggested pro-proliferative effects of endothelins; however, such models poorly recapitulate the *in vivo* glioma context. In this study, we comprehensively revisited endothelin signaling - covering receptor expression, regulation, downstream pathways, and cellular responses-using eleven serum-free, patient-derived glioma lines (glioblastomas, IDH-wt and IDH-mutant oligodendrogliomas and astrocytomas), along with primary tumor samples. Multi-omics and electrophysiological analyses revealed EDNRB as the predominant receptor, enriched in astrocyte-like cells, upregulated by BMPs or growth factor withdrawal, and downregulated by interferons, IL-6 cytokines, endothelins, and Hippo/YAP activation. In contrast, EDNRA was expressed by a perivascular tumor subpopulation and induced by Notch signaling in glioblastomas but not in IDH1-mutant cells. Functionally, endothelins reduced proliferation across all models while promoting migration and PMT. Mechanistically, EDNRB activation increased intracellular Ca²⁺ and activated ERK, STAT3, and apamin-sensitive SK2/SK3 potassium channels. These findings identify endothelin signaling as an important regulator of glioma cell plasticity and behavior.

**Highlights:**

- EDNRB is the predominant endothelin receptor expressed in glioma cells, with a small subset of tumor cells expressing EDNRA in close proximity to blood vessels
- Endothelin signaling reduces proliferation while promoting cell migration and Proneural-to-Mesenchymal transition
- Endothelin activates downstream Ca^2+^, K^+^, ERK, and STAT3 signaling pathways
- EDNRB expression is both positively and negatively regulated by inflammatory cytokines and the Hippo/YAP1 pathway, whereas EDNRA is upregulated by Notch signaling and hypoxia

## Introduction

Diffuse gliomas are the most common type of malignant primary tumors of the central nervous system and remain incurable [1], [2]. Mounting evidence suggests that gliomagenesis originates in adult brain neural stem cells or glial progenitors, which acquire oncogenic alterations and escape normal differentiation cues [3]. Clinically, diffuse gliomas are divided into three major entities: glioblastomas (GB)-the most aggressive form - and lower-grade astrocytomas and oligodendrogliomas [4], [5], [6]. The latter two are distinguished by recurrent mutations in the IDH1 gene, whose aberrant enzymatic activity rewires cellular metabolism and reshapes the epigenome, ultimately driving defective lineage commitment and tumor progression [7]. Single-cell RNA sequencing has revealed profound intra-tumoral heterogeneity across diffuse gliomas [8]. In IDH-mutant astrocytomas and oligodendrogliomas, tumor cells transition among astrocyte-like, oligodendrocyte-like, and neural stem cell-like states [9], [10]. The signaling pathways regulating these lineage identities are beginning to be elucidated. In lower-grade gliomas, we and others have shown that Notch and BMP signaling drive cells toward a quiescent or slowly proliferative, astrocytic-like phenotype [11], [12]. Of note, glioblastomas contain an additional mesenchymal-like population whose abundance varies between tumors [13], [14]. The relative proportions of oligodendrocyte-like and mesenchymal-like cells contribute to the classification of GB into proneural and mesenchymal subtypes, respectively [15]. Frequent proneural-to-mesenchymal transitions (PMTs) are observed, particularly upon recurrence, and are promoted by therapeutic stress (irradiation, temozolomide), hypoxia, inflammation, and neuronal activity [16]. This transition toward a mesenchymal-like state is orchestrated by key transcription factors-including STAT3, C/EBPβ, NF-κB, Notch, and YAP/TAZ-and can be initiated by pro-inflammatory cytokines such as oncostatin M (OSM) and leukemia inhibitory factor (LIF), which could activate the STAT3 pathway and are primarily secreted by immune cells in the tumor microenvironment (TME)[17], [18]. Beyond supplying oxygen and nutrients to cancer cells, vascular cells in the TME also play an essential role in shaping tumor cell identity [19], [20]. They supply Notch ligands that activate Notch signaling in glioma cells and can engage the Hippo/YAP1 pathway, both of which profoundly influence cellular phenotype and behavior [19], [21]. Blood vessels additionally produce bone morphogenetic proteins (BMPs), which likely contribute further to the modulation of lineage states in glioma [22], [23], [24].

Another vascularly-linked pathway of interest is endothelin signaling. Endothelins (ET-1, ET-2, ET-3) are cytokine-like peptides synthesized as ∼212-amino acid preproendothelins that undergo sequential proteolytic processing to yield the mature, bioactive peptides [25]. They signal through two G-protein-coupled receptors, the endothelin receptor type A (EDNRA) and B (EDNRB) [25]. In the brain, endothelial cells are the predominant source of ET-1, but under pathological or stress conditions reactive astrocytes, activated microglia and subsets of neurons also produce ET-1. ET-2 appears chiefly astroglial, whereas ET-3 is enriched in choroid-plexus and circumventricular neurons [26]. Beyond their canonical role in regulating vascular tone, endothelins have also emerged as context-dependent modulators of proliferation, migration and differentiation in astrocytes, neural stem cells and multiple cancers [26], [27].

The presence and impact of endothelins on gliomas have been explored in several studies [28], [29], [30], [31], generally reporting pro-proliferative effects [32], [33], [34]. However, these studies mainly used serum-cultured glioma cell lines, such as rat C6 and human U87, conditions that profoundly alter glioma phenotypes and activate multiple signaling pathways, thereby failing to accurately recapitulate the molecular and functional properties of gliomas [35], [36]. To address this limitation, we undertook a comprehensive reassessment of endothelin signaling in gliomas using a panel of eleven well-characterized, serum-free patient-derived cell lines representing glioblastomas, IDH1-wt and IDH1-mutant oligodendrogliomas and astrocytomas, complemented by analyses of primary tumor specimens. Through an integrative approach combining multi-omics and electrophysiological assays, we re-evaluate the expression and regulation of EDNRA and EDNRB, their downstream signaling pathways, and the functional responses of glioma cells to endothelin stimulation.

## Material and Methods

### Statement on Language Editing Assistance

The writing of this manuscript was refined with the assistance of ChatGPT (OpenAI), which was used to improve grammar, clarity, and style without altering the scientific content.

### Reagents, QPCR primers and Antibodies

References for all products used in this study are provided in the supplementary **Table S1**. A list of QPCR primers is available in **Table S5** and a list of primary antibodies in **Table S6**.

### Patient Samples

Tumor samples were obtained from the Neurology fresh tissue collection of the Montpellier University Hospital Biological Resources Center (Gui de Chauliac; Project No. RECH/P722/1-5) with informed patient consent. Glioma grading was performed by an experienced neuropathologist (Prof. V. Rigau) according to the 2021 WHO classification [4], based on both molecular alteration (IDH1, ATRX, P53, 1p/19q) and histopathological criteria (See **Table S2** for information of patient specimens included in this study). The tumor microarrays (TMAs) used in **Figure S10** are the same as described previously [37], [38]. These TMAs, generated from human brain tumors collected retrospectively at Uppsala University Hospital, include low- and high-grade gliomas as well as non-malignant brain tissue. Each TMA contains duplicate 1 mm tissue cores representing characteristic regions of the tumors, as summarized in **Table S3**. EDNRA abundance was assessed by immunohistochemistry-based protein profiling using an anti-EDNRA antibody (AER-001, Alomone Labs).

### Glioma cell lines

Astrocytoma cell lines (LGG85, LGG275, LGG336, LGG349) were generated in-house from explant cultures of surgical resections performed at Montpellier University Hospital (CHU Gui de Chauliac), following the procedures previously described [39]. The characterization of IDH1-mutant LGG275 and LGG85 has been reported [11], [40]. Detailed molecular characterization of LGG336 (IDH1-mutant) and LGG349 (IDH1-wild type) will be reported elsewhere (Garcia et al., *in preparation*). Oligodendroglioma cell lines included BT138, BT237 [41], BT054 and BT088[42], [43]. Glioblastoma stem-like cell lines (Gb4, Gb5, Gb7, Gb21) were established and characterized as described previously [44], [45]. A complete list of cell lines and key features is provided in **Table S4**. U87 cell line (RRID: CVCL_0022) and T98G glioblastoma cell lines (RRID: CVCL_0556) (gift from Dr N.Laguette’lab, IGH, Montpellier) were cultured with serum according to the guidelines provided in the ATCC catalog.

Additional materials and methods items are provided in the **Supplementary information file**.

## Results

### EDNRB is the main endothelin receptor expressed and is confined to astrocytic-like tumoral cell population in diffuse gliomas

We initiated our analysis of *EDNRA* and *EDNRB* expression in diffuse gliomas by performing qPCR in 14 primary diffuse-glioma resections spanning from low- to high-grade lesions (**Fig. 1A**). *EDNRB* mRNA was detectable in 13 of 14 tumors and, on average, was markedly more abundant than *EDNRA*. Although *EDNRB* levels varied widely between patients, the receptor was present in every histological subtype and grade examined; astrocytoma, oligodendroglioma, and glioblastoma. This predominance of *EDNRB* over *EDNRA* was corroborated by three independent public datasets (TCGA, REMBRANDT, and CGGA, **Fig S1A-D**). In TCGA, both endothelin receptors were expressed more strongly in astrocytomas than in oligodendrogliomas (**Fig. S1D**). Importantly, *EDNRB* expression declined with increasing tumor grade and fell further at recurrence (**Fig. S1B, C**), whereas *EDNRA* showed the opposite pattern, being enriched in high-grade tumors (**Fig. S1B**). Within glioblastoma, *EDNRA* was highest in the mesenchymal subtype, while *EDNRB* predominated in the classical and proneural subtype (**Fig. S1D**). To determine whether the mRNA differences translate at the protein level, we quantified EDNRA and EDNRB proteins in 9 additional samples that covered the main histological glioma subtypes and grades (**Fig. 1B**). Antibody specificity was verified in CHO cells engineered to overexpress either receptor (**Fig. 1B, S4B**). Although the small cohort size precluded robust statistics, both receptors were variably detected across tumors, and EDNRB generally surpassed EDNRA - especially in astrocytomas- mirroring the transcript data. Collectively, these data reveal a dynamic, grade- and subtype-dependent regulation of endothelin receptors in diffuse gliomas, suggesting distinct roles for EDNRA and EDNRB in tumor progression and recurrence.

**Figure 1.**
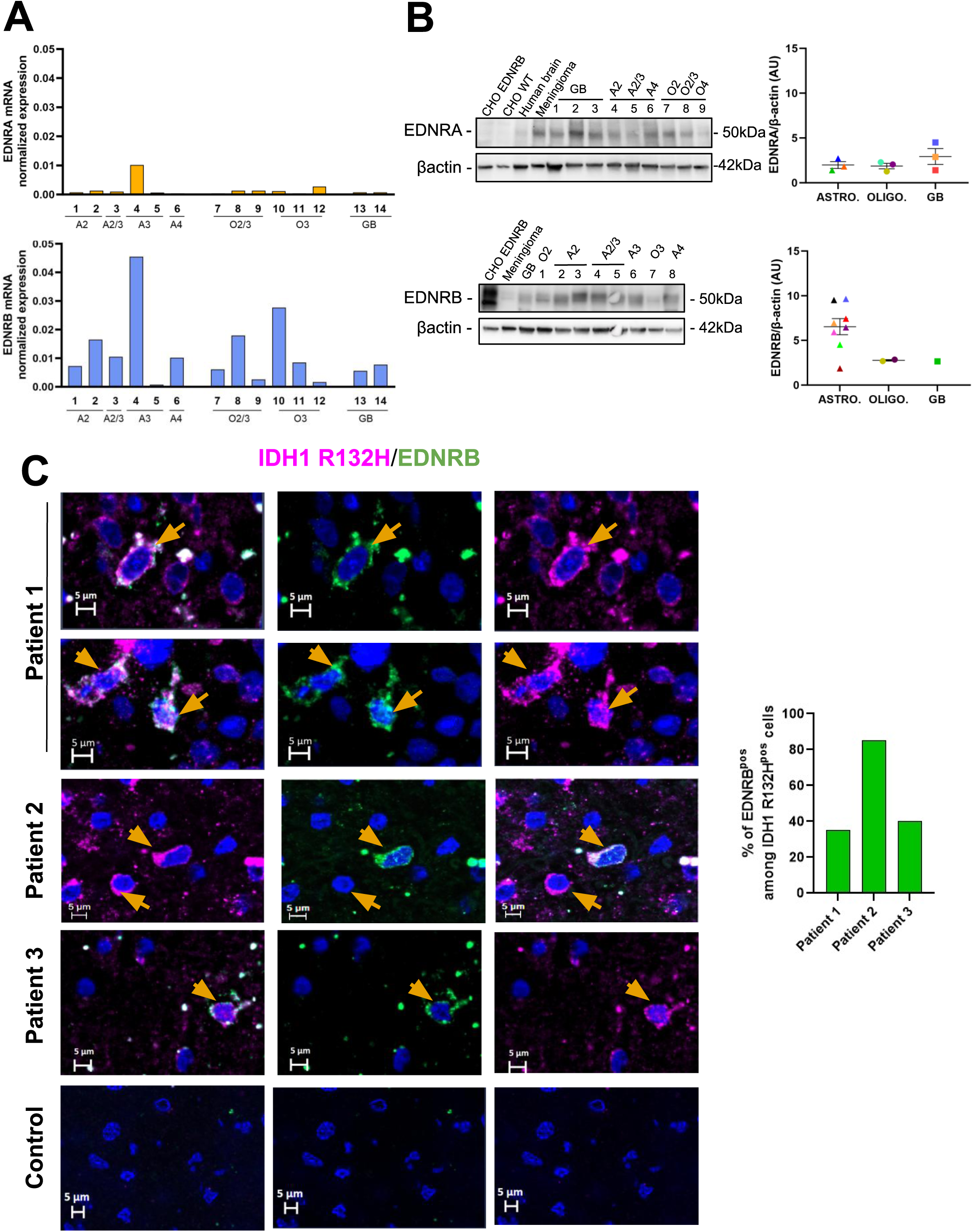
EDNRB is the predominant endothelin receptor expressed in gliomas. **A.** RT-qPCR quantification of *EDNRA* (*top*) and *EDNRB* (*bottom*) mRNA expression in glioma 14 patient samples normalized to *ACTB* gene **B**. (*left)* WB of EDNRA (*top*) and EDNRB (*bottom*) protein expression in 8-9 glioma patient samples. CHO cells (CHO-WT) were used as a negative control; meningioma served as a positive control for EDNRA. CHO cells overexpressing EDNRB (CHO-EDNRB) are used as positive controls for EDNRB antibody and to test specificity of EDNRA antibody. β-actin was used as loading control. (*right):* Quantification of EDNRA and EDNRB levels normalized to β-actin (a.u.). Tumor types: A = Astrocytomas, O = Oligodendrogliomas, GB = Glioblastomas. **C.** Immunofluorescence co-staining of IDH1-R132H and EDNRB in three IDH-mutant diffuse low-grade gliomas. Hoechst: nuclei. Orange arrows: double-positive (IDH1-R132H⁺/EDNRB⁺) tumor cells.

Single-cell transcriptomic data from gliomas revealed that EDNRB is expressed not only in tumor cells but also in non-neoplastic astrocytes (**Fig. S3A**) [46]. To unequivocally confirm its expression in tumor cells, we leveraged IDH1 R132H as a marker in IDH1-mutant gliomas. Employing double labeling of EDNRB and IDH1 R132H in 3 IDH1-mutant patients, we found that a significant number of IDH1 R132H^+^ tumoral cells (25-75%) co-express EDNRB (**Fig. 1C**). Given the now well-established intratumoral cellular heterogeneity of gliomas, we next investigated which tumor cell subtypes express EDNRB in glioblastomas, astrocytomas, and oligodendrogliomas using publicly available single-cell RNA-sequencing datasets. In both glioblastomas and oligodendrogliomas, EDNRB expression appears enriched in cells displaying an astrocyte-like phenotype (**Fig. S2A**). In astrocytomas, correlations between EDNRB and astrocytic or oligodendrocytic markers (**Fig. S2B**) confirm its predominance in tumor cells with an astrocytic signature, mirroring its expression pattern in the normal brain [47]. Notably, these tumoral astrocyte-like cells have been reported to exhibit low proliferative activity in glioblastoma [48]. Consistently, our reanalysis of scRNA-seq data from Neftel et al. revealed that *EDNRB* shows minimal correlation with cyclins and proliferation markers (*mKI67*, *PCNA*), in contrast to oligodendrocyte-lineage genes (*OLIG1*, *OLIG2*, *ASCL1*), which display stronger associations, reflecting their enrichment in actively dividing cell populations (**Fig. S3B**)[13]. To validate at the protein level the preferential expression of EDNRB in astrocyte-like cells, we conducted co-labeling experiments in one astrocytoma and one oligodendroglioma sample, using two markers of astrocytic and oligodendrocytic lineages, namely APOE and OLIG2, respectively. As shown in Figures **S2C** and **S2D**, the vast majority of EDNRB*⁺* cells co-expressed APOE, whereas only a few co-expressed OLIG2, confirming the astrocytic identity of EDNRB-expressing cells.

### Expression of endothelin receptors in glioma cell lines

To identify a suitable *in vitro* model to study endothelin signaling, we performed bulk RNA sequencing on a panel of 11 patient-derived glioma cell lines cultured in a serum-free medium. This panel included three IDH1-mutant lines (LGG275, LGG336, LGG85) derived from low- and high-grade astrocytomas, a high-grade IDH1-wt astrocytoma line (LGG349) [11], [39], [40]. It also comprised four high-grade oligodendroglioma lines (BT054, BT088, BT138, BT237) [41], [42], and previously characterized three glioblastoma cancer stem cell lines (Gb4, Gb7, Gb21) [44], [45]. To assess regulation by growth factors, we also repeated RNA-seq after a 4-day growth-factor withdrawal, a condition promoting lineage commitment and proliferation reduction. *EDNRB* was more strongly expressed than *EDNRA* in most cell lines but absent in four of them (LGG85, LGG349, BT138 and BT088). In the seven positive cell lines, *EDNRB* transcripts rose after growth-factor withdrawal whereas *EDNRA* remained undetectable (**Fig. 2A**); qPCR confirmed these results **(Fig. S4A**). Protein validation mirrored the RNA profile: WBs showed heterogeneous, but concordant EDNRB expression that increased after growth-factor removal (**Fig. 2B**). EDNRB migrated as two to three bands, suggesting alternative splicing and/or post-translational modifications. Consistent with RNA, EDNRA protein was not detected in any of the four cell lines examined (**Fig. S4B**). Then, to assess surface expression of the EDNRB receptor, we performed flow cytometry and IF on non-permeabilized live LGG275 cells using two independent antibodies targeting the extracellular domain of the receptor. Both approaches confirmed membrane localization of EDNRB and confirmed its upregulation following growth factor withdrawal (**Fig. S4C, D**).

**Figure 2.**
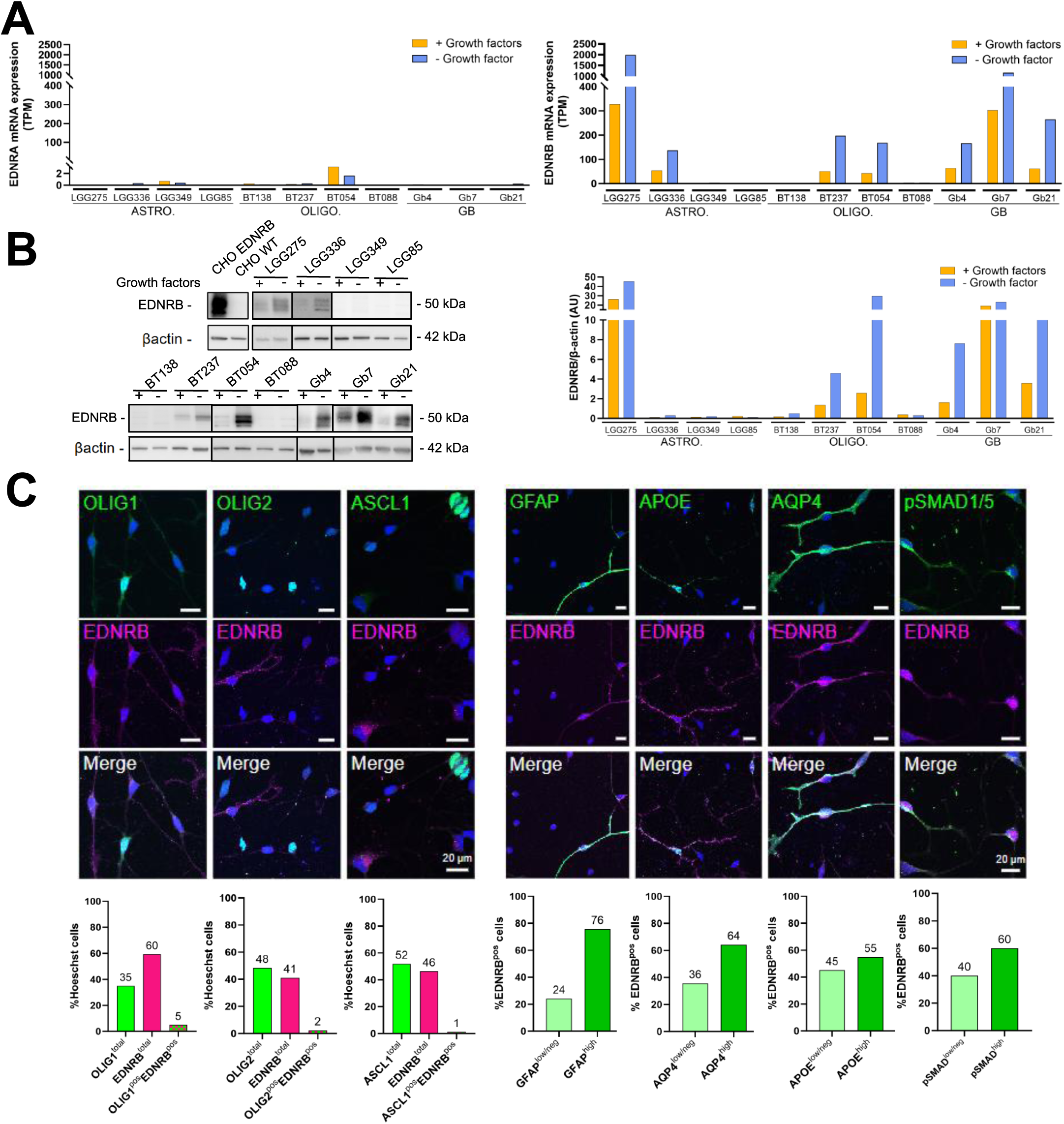
EDNRB dominates endothelin receptor expression in glioma cell lines. **A.** RNA-seq analysis of *EDNRA* and *EDNRB* (TPM) in 11 glioma cell lines cultured ±growth factors (GFs): astrocytomas (LGG275, LGG336, LGG85, LGG349), oligodendrogliomas (BT138, BT237, BT054, BT088), and IDH–wild-type glioblastomas (Gb4, Gb7, Gb21). **B.** Western blot of EDNRB in the same lines ±GFs, with CHO-WT (negative) and CHO-EDNRB (positive) controls; β-actin: loading control. Right: densitometric quantification (a.u.). **C.** Immunofluorescence of LGG275 cells showing EDNRB⁺ cells lack oligodendroglial (OLIG1, OLIG2) and neural progenitor (ASCL1) markers but variably express astrocytic markers (GFAP, APOE, AQP4, pSMAD1/5). Nuclei: Hoechst.

As we found that EDNRB was mainly expressed by astrocytic-like cells in patients (**Fig. S2A-D**), we tested whether this pattern was recapitulated *in vitro* using LGG275 cells [11] which showed higher *EDNRB* RNA expression than all other cell lines analyzed (**Fig. 2A**). After growth factor withdrawal, LGG275 cells display 2 distinct populations expressing either oligodendrocyte-lineage transcription factors (OLIG1, OLIG2, ASCL1) or astrocyte-lineage markers (GFAP, APOE, AQP4, pSMAD1/5) (**Fig. 2C**). Double immunostainings showed EDNRB was largely absent from oligodendrocytic cells but co-expressed with astrocytic markers, indicating its restriction to astrocytic-like cells in LGG275. scRNA-seq analysis of LGG275 further confirmed *EDNRB* expression in cells with an astrocyte-like transcriptional profile, as observed in patients (**Fig. S3D)**. This analysis also revealed that *EDNRB*-positive cells in LGG275 exhibit reduced expression of proliferation markers, including *PCNA* and *MKI67*, suggesting a lower proliferative state within this subpopulation (**Fig. S3C)**.

### Endothelins reduce glioma cell proliferation and promote cell motility

To evaluate how endothelin-receptor stimulation influences cell growth, we treated ten glioma cell lines with either ET-1 or ET-3 for 10–15 days, then quantified total cell numbers (**Fig. 3A-B**). Six of the treated lines exhibited a moderate yet significant reduction (≈10–20 %) in cell number. The four lines that failed to respond were those lacking EDNRB expression (**Fig. 2A-B**) underscoring its role in the growth-inhibitory effect. Focusing on the LGG275 line, dose–response assays with ET-1/ET-3 showed growth inhibition from 4 nM (**Fig. 3C**). The selective EDNRB agonist IRL-1620 reproduced this effect, confirming EDNRB receptor involvement (**Fig. 3D**). To assess whether the reduction was due to decreased proliferation or increased apoptosis, we performed EdU incorporation assays and apoptosis detection. Both ET-1 and IRL-1620 significantly reduced the proportion of EdU-positive cells (**Fig. 3E)**, whereas no significant change in apoptosis was observed (**Fig. S5A**). RNA-seq (*see below*) further supported the anti-proliferative effect of endothelin signaling showing a 15% decrease in mKI67 and PCNA transcript levels (**Fig. 3F**).

**Figure 3.**
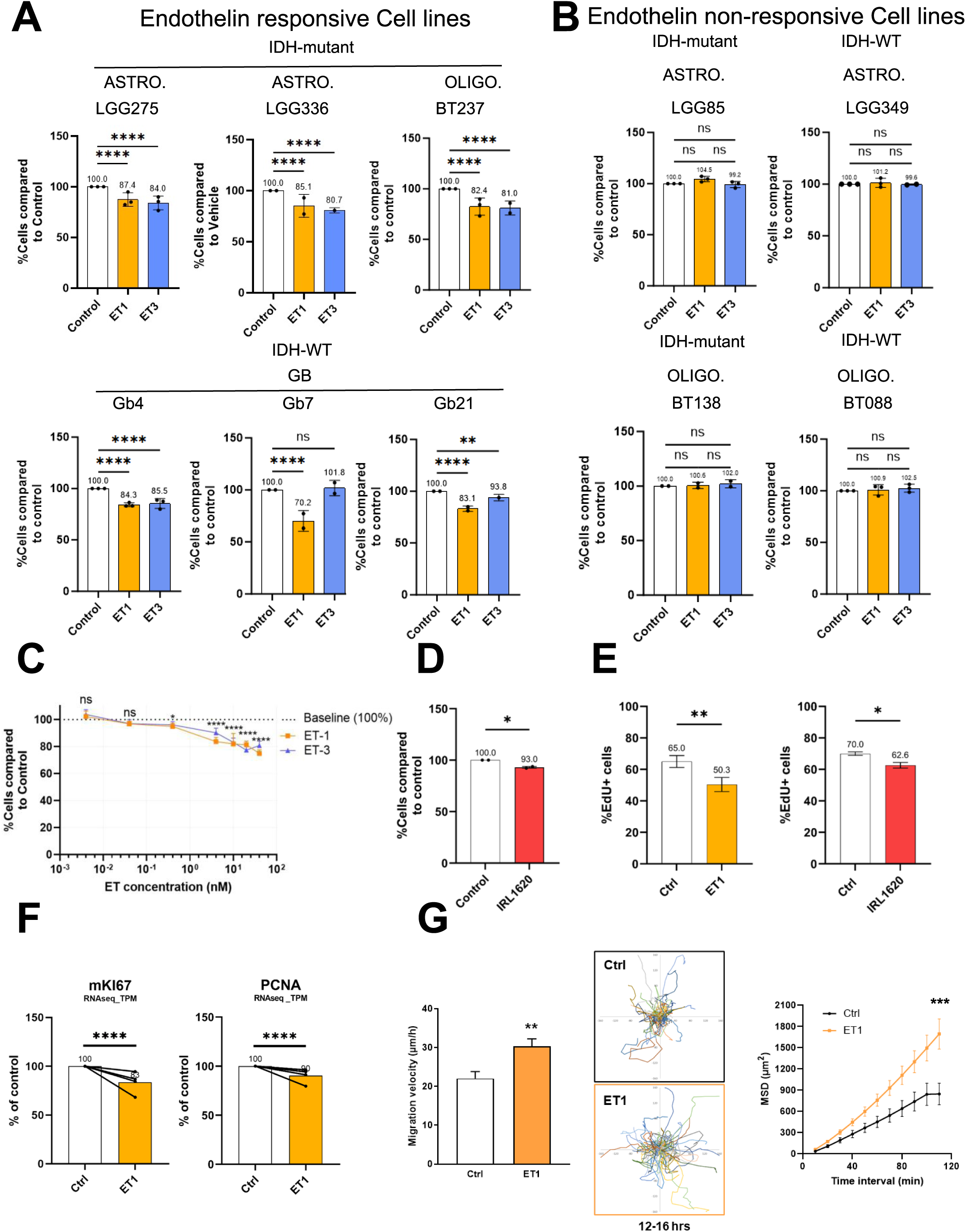
Endothelins reduce proliferation in a subset of diffuse glioma cell lines and promote migration. **A–B.** Effect of ET-1 and ET-3 (4 nM) on glioma cell growth. *(A)* In responsive lines, both ligands significantly reduced cell numbers relative to vehicle (100%). *(B)* Non-responsive lines showed no significant change. Data are mean ± SD from three independent experiments (≥12 wells/condition); one-way ANOVA with Dunnett’s post-hoc test. **** p <0.0001; *** p <0.001; ** p <0.01; * p <0.05; ns: not significant. **C.** Dose–response in LGG275 cells showing significant growth inhibition by ET-1 and ET-3 from 0.5 nM. Mean ± SD; ≥12 wells/condition; ANOVA with Dunnett’s post-hoc test. **D.** EDNRB agonist IRL1620 (1 µM) reduces LGG275 growth (mean ± SD; ≥12 wells; n=2). **E.** Percentage of EdU⁺ proliferating LGG275 cells after ET-1 (10 nM) or IRL1620 (1 µM) treatment. ET-1: n=5; IRL1620: n=2 with 6 technical replicates; unpaired t-test (ET-1) or ANOVA with Dunnett’s test (IRL1620). ** p <0.01; * p <0.05. **F.** Decreased *MKI67* (left) and *PCNA* (right) mRNA (TPM) in LGG275 after ET-1 (10 nM) (n=4; Bootstratio test). **** p <0.0001.**G.** Migration assays in LGG275 cells with or without ET-1 (10 nM). *(Left)* Mean migration velocity over 12–16 h post-treatment (**p <0.01, t-test). *(Middle)* Plot-to-origin trajectories showing more cells migrating farther. *(Right)* Mean square displacement (MSD) over 10 min intervals from 12–16 h, indicating greater area explored with ET-1 (***p <0.001; two-way ANOVA).

Finally, as a reduction in proliferation can coincide with greater migratory capacity, a feature of the “go-or-grow” paradigm [49], we asked whether ET-1 promotes cell motility. We therefore quantified the motility of LGG275 cells by time-lapse microscopy, as previously described [50]. On a laminin-coated substrate, LGG275 cells exhibited a mean migration velocity of 22.0 µm/h. Addition of ET-1 (10 nM) significantly increased migration speed (30.3 µm/h) and territory explored, as shown by cumulative traces and mean square displacement (MSD) (**Fig. 3G**). Experiments conducted at two different cell densities showed no difference in either basal migration velocity or the increase in migration velocity induced by endothelin treatment (*data not shown*). We then assessed the onset of increased migration in endothelin-treated LGG275 cells by measuring 4-hour intervals over a 2–22-hour period. Figure **S5B** shows that migration significantly increased 10 hours after ET-1 addition (+139% in migration velocity) and remained elevated until 22 hours (+173%). Similarly, ET-1 enhanced the spatial exploration of the cells, as highlighted by the plot-to-origin graphs and MSD data (**Fig. S5C, D**). Altogether, our results demonstrate that ET-1 stimulates the motility of LGG275 cells.

### Endothelins promote a proneural-to-mesenchymal transition in glioma cells

To investigate how endothelins affects glioma cells at the molecular level, we performed RNA seq on ET-1 treated LGG275 (n = 3; **Fig. 4**), LGG336, BT237 and Gb4, 7, 21 lines (n=1 each, **Fig. S6**). In LGG275, the volcano plot (**Fig. 4A, Table S7**) shows 50 significantly upregulated and 15 downregulated genes (log₂FC > |1|, q <0.05). Gene Set Enrichment Analysis (GSEA) revealed a reduction in cell cycle–related programs, notably E2F targets and the G2/M checkpoint, along with upregulation of migration-associated genes (**Fig. S5E**, **Table S8**), consistent with our proliferation and migration functional assays. Unexpectedly, GSEA using the Verhaak list of transcriptional signatures that define the classical, proneural, mesenchymal, and neural subtypes of glioblastoma revealed that ET-1 treatment induces a proneural-to-mesenchymal transition (PMT) **(Fig. 4B, Table S9**) [51]. Notably, six well-characterized mesenchymal-associated genes were upregulated following ET-1 treatment: *CCN1/CYR61* [52], *CCN2/CTGF*, *CD44*, *SERPINE1*[16], [17], and *CAV1* [53]. GSEA further showed that ET-1 activates stress- and inflammation-related transcriptional programs, with enrichment of gene sets linked to hypoxia, apoptosis, and reactive oxygen species (ROS), and downregulation of UV-response genes (**Fig. 4C; Table S8**). The enrichment of the epithelial–mesenchymal transition (EMT) hallmark found by GSEA likely reflects a PMT rather than a true EMT, given the non-epithelial origin of glioma cells.

**Figure 4.**
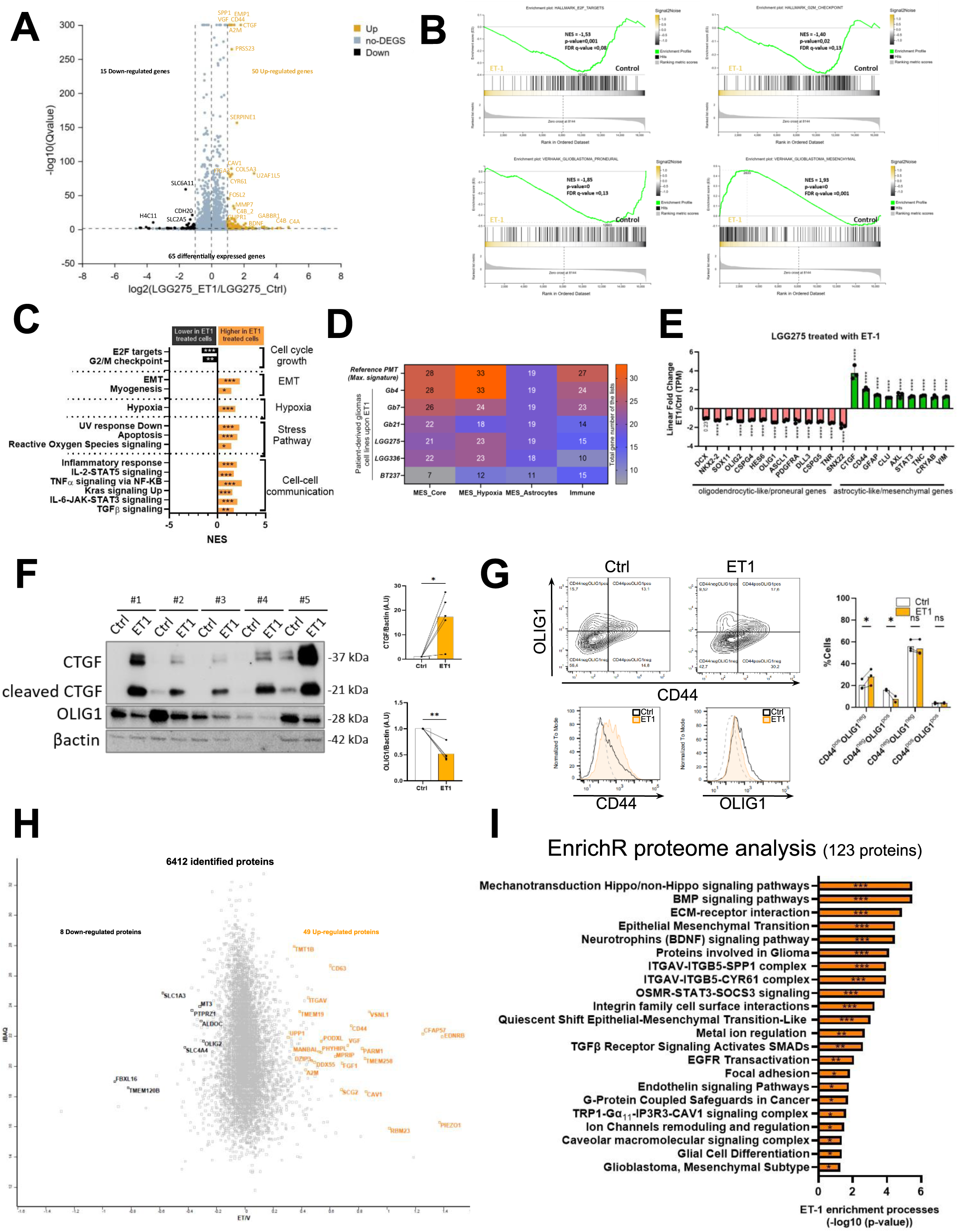
Endothelins promote a proneural-to-mesenchymal transition (PMT) in glioma cell lines. **A.** Volcano plot of differentially expressed genes (DEGs) in LGG275 cells treated with ET-1 (10 nM) versus vehicle, based on RNA-seq (TPM). X-axis: log₂ fold change; Y-axis: –log₁₀(FDR q-value). **B.** Gene Set Enrichment Analysis (GSEA) of ET-1–treated LGG275 cells using MSigDB Hallmark and Curated C2 (CGP) collections. *(Top)* Downregulation of cell cycle–related gene sets (E2F targets, G2/M checkpoint). *(Bottom)* Decreased enrichment of Proneural and increased enrichment of Mesenchymal signatures (Verhaak_Glioblastoma). NES, nominal p-value, and FDR q-value are shown. **C.** Summary of significantly enriched GSEA Hallmark pathways (MSigDB) (FDR q <0.25) in ET-1–treated LGG275 cells. Pathways upregulated are shown in orange; downregulated, in black. *** p <0.001; ** p <0.01; * p <0.05 (nominal p-value). **D.** Heatmap showing overlap between ET-1–upregulated genes in glioma cell lines and mesenchymal gene signatures from Chanoch-Myers et al. [10] (MES_core, MES_Hypoxia, MES_Astrocyte, Immune). Top row: total genes per signature; rows below: number of overlapping genes in ET-1–treated lines. Color intensity and values indicate absolute counts, with higher counts reflecting a stronger shift toward the corresponding mesenchymal subtype. **E.** Fold change in lineage marker expression from RNA-seq (TPM) in LGG275 cells treated with ET-1, showing oligodendrocyte progenitor markers (purple) and astrocytic markers (green). Mean ± SD; n = 3; q-values from DESeq2. **F.** (*Left*) Western blot of mesenchymal/astrocytic-like marker CTGF (full-length and cleaved) and proneural/oligodendrocytic-like marker OLIG1 in LGG275 cells ±ET-1. β-actin: loading control. (*Righ*t) Quantification (A.U.), mean ± SD; n = 5; paired t-test. * p <0.05; ** p <0.01. **G.** Flow cytometry of CD44 and OLIG1 in LGG275 cells ±ET-1 (no GFs). *Top*: density plots; bottom: overlay histograms for CD44 and OLIG1. *Right*: quantification of CD44/OLIG1 subpopulations (mean ± SD; n = 3); two-way ANOVA with Šídák’s test. * p <0.05; ns: not significant. **H.** Scatter plot of proteomic changes in LGG275 cells ±ET-1 by TMT-Mass spectrometry. X-axis: ET-1/vehicle ratio; Y-axis: iBAQ intensity. Shown are proteins significant by Fisher’s exact test (p <0.05) and detected in ≥3/5 experiments with consistent direction. **I.** EnrichR analysis of 123 ET-1–regulated proteins, showing top enriched pathways ranked by –log₁₀(p-value). *** p <0.001; ** p <0.01; * p <0.05.

This shift toward a more mesenchymal phenotype is not limited to LGG275. GSEA of RNA-seq from LGG336 and the Gb4, Gb7, and Gb21 lines also revealed an EMT signature (**Fig. S6 B-C, Tables S10-19**). BT237 was the sole exception, indicating that oligodendroglioma cells may be less susceptible to ET-1-induced PMT transition. As observed in LGG275, ET-1 also induced the expression of inflammation-related genes and activated the IL6-JAK-STAT3 signaling pathway in most lines.

Whereas mesenchymal (MES) programs are recurrently reported in gliomas, yet their exact composition varies across studies. A recent meta-analysis of ten published MES signatures distilled a 28-gene “MES-core” that appears in at least three independent datasets [14]. In our RNA-seq dataset, ET-1 treatment upregulated the vast majority of these core genes – 22 of 28 genes -in five of the six patient-derived lines examined (LGG275, LGG336, Gb4, Gb7, Gb21), whereas the oligodendroglioma line BT237 displayed only a marginal response (**Fig. 4D, Table S20**). The same study further resolved three recurrent MES sub-states: i/MES-Hyp, dominated by hypoxia/glycolysis genes; ii/ MES-Ast, enriched for astrocytic markers and antigen-presentation genes; iii/ An intermediate state that co-expresses elements of both programs. Cross-referencing these sub-state signatures with our ET-1 dataset revealed a strong concordance with the hypoxia module and, to a lesser extent, enrichment of selected astrocytic genes (*CLU, GFAP, GJA1, HOPX, SPARC*). Additionally, of the seven gene sets linked to the mesenchymal (MES) phenotype in this meta-analysis, our data showed the strongest overlap with the immune/MHC cluster in the Gb4 and Gb7 cell lines, consistent with reports that this MES cluster distinguishes glioblastoma from lower-grade gliomas [14]. Collectively, these results indicate that ET-1 robustly drives a mesenchymal transition dominated by the MES-Hyp type, while partially engaging astrocytic and immune-associated pathways.

Given that acquisition of mesenchymal features is typically associated with repression of proneural/oligodendrocyte progenitor cell markers, we next extracted from our RNA seq dataset the expression of 13 canonical OPC genes across our six glioma cell lines. Indeed, ET-1 treatment downregulated most OPC markers in five lines, with BT237 (oligodendroglioma) as the exception (**Fig. 4E, S6B–C**). In contrast, a curated panel of astrocytic and MES-associated genes (*CTGF, CD44, GFAP, CLU, AXL, STAT3, TNC*, *CRYAB* and *VIM*) was upregulated in the same five lines, again with minimal or no response in BT237 (**Table S21**). Protein-level analyses confirmed these transcriptional shifts. We first focused on OLIG1, a key OPC transcription factor, and *CCN2/CTGF* a prototypical mesenchymal matrix protein induced by TGF-β, hypoxia, and inflammation [54]. LGG275 cells showed strong induction of both full-length and cleaved CTGF isoforms accompanied by a decrease in OLIG1 levels in WB (**Fig. 4F**). Flow cytometry further supported this phenotypic shift: the proportion of CD44^pos^/OLIG1^neg^ cells increased, while CD44^neg^/OLIG1^pos^ OPC-like cells decreased following ET-1 treatment (**Fig. 4G**), consistent with a proneural-to-mesenchymal transition. To assess global protein-level changes, we performed quantitative mass spectrometry in LGG275 upon ET-1 (n=3). ET-1 significantly upregulated 49 proteins and downregulated 8 (ratio ET/ctrl significant in at least 3 out of 5 replicates) (**Fig. 4H**, **Tables S22-23**). Notably, two proteins characteristic of the proneural state (OLIG2 and PTPRZ1) were downregulated, whereas two canonical MES markers (CAV1 and CD44) were upregulated. Enrichment analysis of upregulated proteins using EnrichR [55] revealed significant enrichment in pathways related to epithelial–mesenchymal transition (EMT), STAT3 signaling, and integrin–extracellular matrix interactions reminiscent of PMT (**Fig. 4I, S7A-B, Tables S24-25**).

### Endothelins stimulate Ca2+, ERK and STAT3 signaling in LGG275

Mesenchymal transition in gliomas is largely driven by STAT3 signaling, with ERK and calcium pathways also implicated [56], [57], [58]. Our RNA-seq and proteomic data suggested activation of these pathways, leading us to test whether endothelins stimulate Ca²⁺, ERK, and STAT3 signaling in LGG275 cells. We first examined calcium signaling. As EDNRB canonically couples to Gαq/11–PLC–IP3, we measured IP1 accumulation, a downstream metabolite of IP3, using the FRET-based HTFR assay. ET-1 or ET-3 (40 nM) significantly increased IP1, with stronger effects under growth-factor–deprived conditions (**Fig. 5A)**, consistent with the previously noted EDNRB upregulation (**Fig. 2A-B**). The EDNRB agonist IRL-1620 reproduced this effect, which was abolished by the competitive antagonist BQ788, confirming specificity (**Fig. 5B**). To directly assess Ca²⁺ mobilization, we used Cal-520-AM (**Fig. 5C-D**) and Fura-2-AM (**Fig. 5E**), both showing robust ET-1–induced Ca²⁺ increases, further enhanced without growth factors, again sensitive to agonist and antagonist. Similar EDNRB-dependent Ca²⁺ responses to ET-1/ET-3 were observed in BT237 cells (**Fig. S8A**).

**Figure 5.**
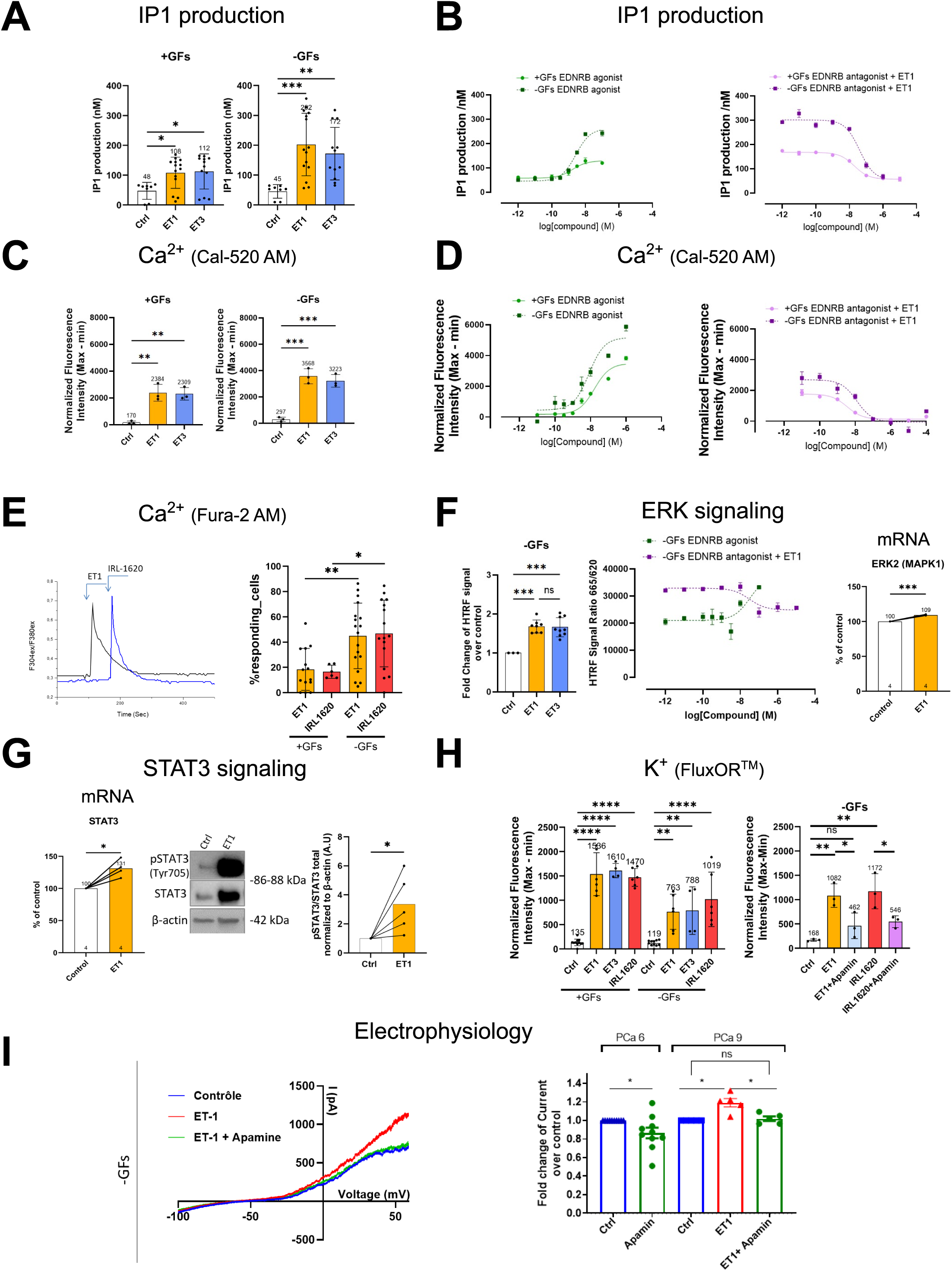
Endothelin signaling stimulates Ca²⁺, K⁺, ERK, and STAT3 pathways in LGG275 cells. **A.** IP₁ production measured by HTRF in LGG275 cells cultured with (+GFs) or without (–GFs) growth factors after ET-1 or ET-3 treatment. Mean ± SD from n = 5 independent experiments; unpaired t-test. *** p <0.001; * p <0.05. **B.** IP₁ production in response to the EDNRB agonist IRL-1620 (left) or antagonist BQ-788 plus EC₈₀ ET-1 (4 nM) (right) under ±GF conditions. Non-linear regression dose– response curves from one representative of three experiments. **C.** Intracellular Ca²⁺ flux (Cal-520AM) in control, ET-1, or ET-3 conditions under ±GFs. Mean ± SD; n = 3 independent experiments; one-way ANOVA/Tukey. *** p <0.001; ** p <0.01; * p <0.05. **D.** Ca²⁺ flux (Cal-520AM) in response to IRL-1620 (left) or BQ-788 plus ET-1 (right) under ±GFs. Non-linear regression curves from one representative of three experiments. **E.** EDNRB-dependent Ca²⁺ responses measured by Fura-2. *(Left)* Representative traces for ET-1 (40 nM) or IRL-1620 (1 µM). *(Right)* Percentage of responding cells under ±GFs. Mann–Whitney test. ** p <0.01; * p <0.05. **F.** ERK activation. *(Left)* pERK1/2 quantified by HTRF 5 min after ET-1 or ET-3 (40 nM) in –GF conditions (n = 3). *(Middle)* pERK1/2 dose– responses to IRL-1620 or BQ-788 plus ET-1 (4 nM) in –GF conditions. Mean ± SD; ANOVA/Tukey. *** p <0.001; ns: not significant. Representative result from three independent experiments. *(Right) MAPK1* (ERK2) mRNA levels in ET-1–treated versus control cells (RNA-seq, TPM; n = 4 independent experiments). **G.** STAT3 activation. *(Left) STAT3* mRNA (TPM) expression after ET-1 treatment (n = 4 independent experiments). *(Middle)* WB of pSTAT3 (Tyr705), total STAT3, and β-actin. *(Right)* Quantification of pSTAT3 normalized to total STAT3 and β-actin (A.U.), relative to control, from n = 5 independent experiments. Paired t-test. * p <0.05. **H.** K⁺ fluxes (FluxOR™ probe) under ±GFs in response to ET-1, ET-3, or IRL-1620. *(Left)* Normalized fluorescence (Max–Min) from n = 3 independent experiments; ANOVA/Tukey tests. *(Right)* K⁺ fluxes under –GFs induced by ET-1 or IRL-1620 and blocked by apamin (1 µM) a blocker of SK channels. Multiple unpaired t-tests. **** p <0.0001; *** p <0.001; ** p <0.01; * p <0.05; ns: not significant. **I.** Whole-cell voltage-clamp recordings under –GFs. *(Left)* I–V curves showing ET-1–induced outward K⁺ currents (red) blocked by apamin (green) versus control (blue). *(Right)* Fold change in current amplitude under high (PCa6, 1 µM) or physiological (PCa9, 1 nM) intracellular Ca²⁺. At PCa6, currents were reduced by apamin; at PCa9, ET-1 increased current amplitude, partially blocked by apamin. Mean ± SD; n = 6–7 cells; unpaired t-test. * p <0.05; ns: not significant.

To assess MAPK/ERK activation, we quantified ERK1/2 phosphorylation (Thr202/Tyr204) by HTRF. In growth factor–rich medium, ERK1/2 was near saturation, with no further increase upon markedly increased phosphorylation, an effect reproduced by IRL-1620 and reduced by BQ788, confirming EDNRB dependence **(Fig. 5F).** Additionally, ET-1 upregulated MAPK1 (ERK2) mRNA (RNA seq), corroborating pathway activation. STAT3 signaling was likewise induced: RNA-seq showed upregulation of STAT3 mRNA, and WB confirmed a significant increase in pSTAT3/STAT3 upon endothelin treatment (**Fig. 5G, S8B**). Together, these data demonstrate that endothelins stimulate Ca²⁺, ERK1/2, and STAT3 signaling via EDNRB in glioma cells, consistent with their role in mesenchymal transition.

### Endothelins activates an apamin-sensitive Ca²⁺-regulated K⁺ current in LGG275 cells

We previously showed that LGG275 cells express functional an apamin-sensitive K**⁺** channel (*KCNN3/SK3)*, which are activated by intracellular Ca2+ [11], [40]. RNA seq also revealed the expression of another calcium-activated K**⁺**channel, namely *KCNN2/ SK2.* Both channels are detected at the protein level with variation upon growth factor removal (**Fig. S8C-D**). Given that endothelins strongly elevate intracellular Ca²⁺ and *EDNRB* expression correlates with *SK2/3* in glioma patient datasets (**Fig. S8E**), we hypothesized that SK channels are activated downstream of endothelin signaling. To test this, we used the FluxOR™ probe, which enables real-time monitoring of K⁺ channel activity in living cells. Its functionality was validated with the K⁺ ionophore nigericin, which induced a clear fluorescence response (**Fig. S8F**). A Ca²⁺ ionophore further confirmed that Ca²⁺ influx activates K⁺ channels in LGG275 cells, with partial inhibition by apamin under growth factor–free conditions (**Fig. S8F**). Using this probe, we found that ET-1, ET-3, and IRL-1620 all triggered measurable K⁺ fluxes in both growth factor–rich and –deprived media, a response significantly reduced by apamin, indicating mediation, at least in part, by apamin-sensitive K⁺ channels (**Fig. 5H**). To further confirm functional SK channel activation, we performed whole-cell electrophysiology. Under high intracellular Ca²⁺ (PCa6, 1 µM), cells displayed spontaneous SK-type currents partially blocked by apamin (**Fig. 5I, S8G**). At lower, physiological Ca²⁺ (PCa9, 1 nM), ET-1 application evoked an apamin-sensitive current, demonstrating that ET-1 activates SK channels (**Fig. 5I**). Although pharmacology did not allow us to distinguish SK2 from SK3, these data show that endothelins stimulate apamin-sensitive K⁺ channels in LGG275 cells.

### BMPs, IL-6–related cytokines and endothelins differentially control EDNRB expression

In light of EDNRB’s functional impact on glioma cells, we then investigated the regulation of its expression at both RNA and protein levels. We first observed that treatment of LGG275 cells with ET-1 led to a reduction in both EDNRB mRNA and protein levels, suggesting the presence of an intrinsic negative feedback loop whereby endothelin downregulates its own receptor expression (**Fig. 6A**). Since EDNRB is enriched in astrocyte-like tumor cells, we next tested whether microenvironmental cues promoting astrocytic differentiation affect EDNRB levels. We first treated LGG275 cells with BMP2, which are secreted by glioma cells, reactive astrocytes, and endothelial cells [24]. All three BMPs significantly increased *EDNRB* mRNA, and BMP4 also elevated EDNRB protein (**Fig. 6B**). We next tested other pro-inflammatory cytokines that promote astrocytic fate— LIF, OSM, CNTF and IFN-γ. In sharp contrast to BMPs, each of these cytokines markedly repressed *EDNRB* transcription, and EDNRB protein became almost undetectable after LIF or OSM treatment (**Fig. 6C**). Because LIF and OSM activate STAT3 transcription factor, we transduced LGG275 cells with a constitutively active STAT3 mutant; this alone significantly reduced *EDNRB* mRNA (**Fig. 6D**), implicating STAT3 in the repression.

**Figure 6.**
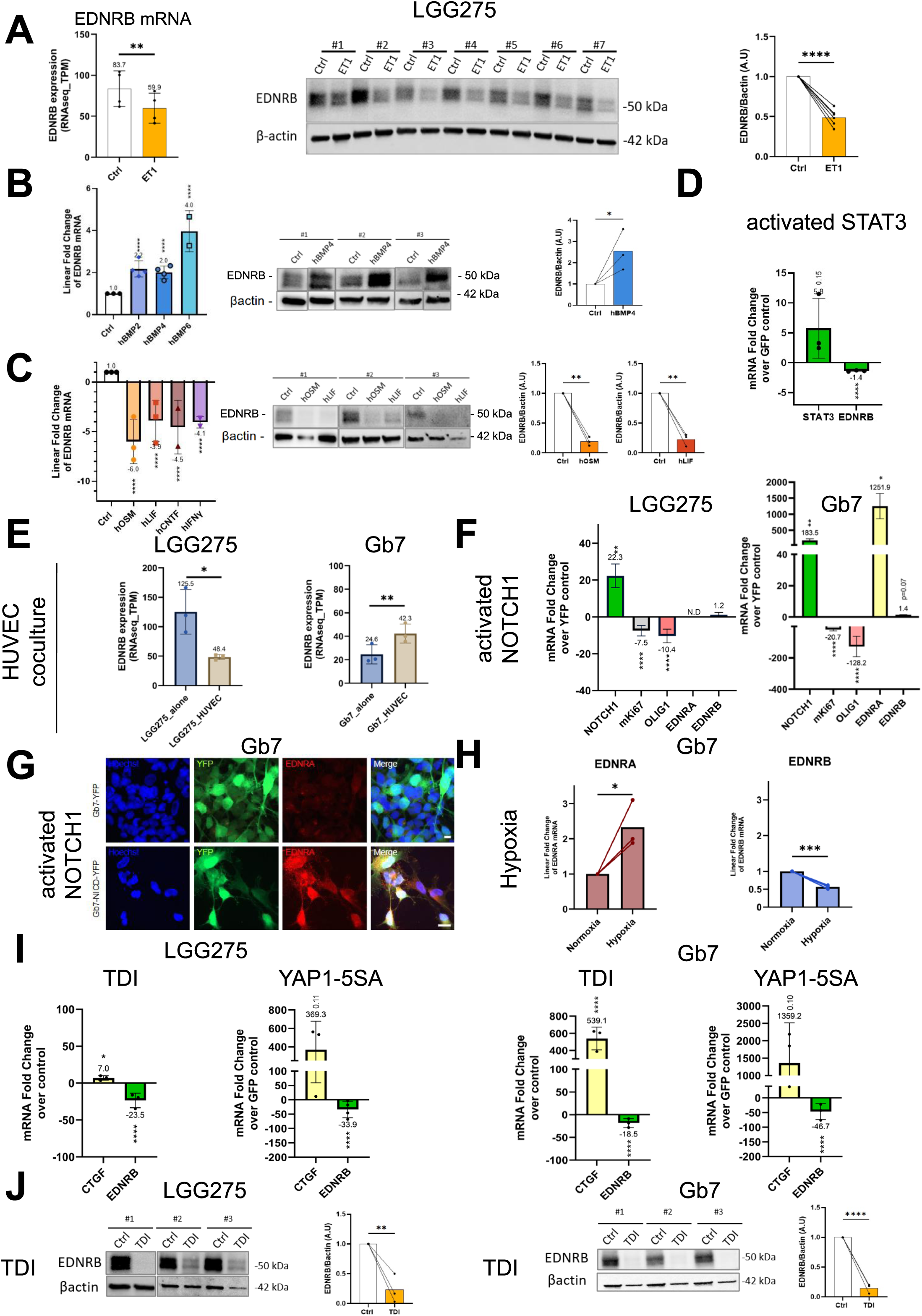
Dynamic regulation of endothelin receptor expression in diffuse glioma cells. **A.** ET-1 reduces *EDNRB* in LGG275 cells. *(Left)* mRNA (RNA-seq, TPM; n=4 independent experiments). *(Middle)* Representative WB (n=7 independent experiments). *(Right)* Quantification (A.U., β-actin–normalized). Unpaired (RNA) or paired t-test (WB), ** p<0.01, ****p<0.0001. **B.** BMPs (10 ng/mL) increase *EDNRB* expression. *(Left)* mRNA (RT-qPCR, n=3 independent experiments). *(Middle)* WB for hBMP4 (n=3 independent experiments). *(Right)* Quantification. Bootstratio (RNA) and paired t-test (WB); ****p<0.0001, *p<0.05. **C.** IL-6 family cytokines (10 ng/mL) (hOSM, hLIF, hCNTF) and IFNγ reduce *EDNRB* in LGG275. *(Left)* mRNA (RT-qPCR, n=3 independent experiments). *(Middle)* WB for hOSM/hLIF (n=3 independent experiments). *(Right)* Quantification. Bootstratio (RNA) or paired t-test (WB); ****p<0.0001, **p<0.01. **D.** Constitutively active STAT3 decreases *EDNRB* mRNA (n=3 independent experiments). Bootstratio; ****p<0.0001. **E.** HUVEC endothelial cells co-culture reduces *EDNRB* in LGG275 and Gb7 cells (RNA-seq, TPM; n=3 independent experiments). Unpaired t-test; *p<0.05, **p<0.01. **F.** Notch activation (NICD-YFP) decreases MKI67 and OLIG1, increases EDNRA in Gb7 but not LGG275, and leaves EDNRB unchanged (RT-qPCR; n=3 independent experiments; Bootstratio; *** p<0.0001; * p<0.05). **G.** Immunofluorescence: NICD (NICD-YFP virus) increases EDNRA⁺ cells in Gb7 cultures. Scale bar: 10 µm. **H.** Hypoxia (1% O₂, 5 days) increases *EDNRA* and decreases *EDNRB* in Gb7 (RT-qPCR; n=3). *p<0.05, ***p<0.001. **I.** Hippo/YAP1 activation by TDI (5 µM) or YAP1-5SA increases *CTGF* and decreases *EDNRB* in LGG275 and Gb7 (RT-qPCR; n=3 independent experiments). Bootstratio; *p<0.05. ****p<0.001 **J.** TDI reduces EDNRB protein in LGG275 and Gb7 (WB; n=3 independent experiments). Paired t-test; **p<0.01, ****p<0001.

These results show that EDNRB expression is enhanced by BMPs but suppressed by STAT3-activating cytokines and ET-1 itself, revealing a dynamic regulatory network that may shape glioma heterogeneity.

### Endothelin receptors expression is controlled by endothelial cells, hypoxia, Notch and Hippo/YAP1 pathways

Blood vessels greatly influence glioma cells by providing ligands that activate Notch1 and Hippo/YAP1 signaling, pathways critical for stem and cancer stem cell maintenance [19], [20], [45], [61]. However, the roles of endothelial cells, Notch1, and Hippo/YAP1 in regulating endothelin receptors remain poorly defined [62]. To address this, as performed previously [44], we co-cultured mCherry^+^ endothelial cells (HUVEC) and IDH1-mutant LGG275 or IDH1-wt. GB Gb7 GFP^+^ cells and then sort GFP^+^ cells and performed RNA-seq (n=3) to assess EDNRA and EDNRB expression. Surprisingly, LGG275 and Gb7 react differentially by downregulating and upregulating EDNRB, respectively, without inducing EDNRA (**Fig. 6E**). To directly test the effect of Notch1 activation, we transduced LGG275 and Gb7 cells with a constitutively active form (NICD) (**Fig. 6F**). As reported previously [11], [45], NICD reduced proliferation (*mKI67*) and the proneural transcription factor *OLIG1*. While EDNRB was unchanged, EDNRA was strongly induced (>1000-fold) in Gb7 but remained undetectable in LGG275. This strong induction in Gb7 was further confirmed by IF (**Fig. 6G)**. Given that hypoxia has been shown to work in a Notch-dependent manner and increases the expression of direct Notch target genes [63], [64], we also assessed the impact of hypoxia on endothelin receptor expression in Gb7 cells. As shown in Figure **6H**, exposure to reduced oxygen levels led to a moderate but significant increase in *EDNRA* expression, accompanied by a concomitant decrease in *EDNRB*. Finally, we explored the impact of Hippo/YAP1 pathway activation both pharmacologically using TDI-011536, an inhibitor of LATS1/2 kinase for YAP1 and by transducing cells with an active form of YAP1 (5SA). Both treatments robustly upregulated CTGF RNA and protein, confirming pathway activation (**Fig. 6I, S9**). Under these conditions, EDNRA remained undetectable (data *not shown*), whereas EDNRB was strongly repressed at RNA and protein levels (**Fig. 6I–J**).

Together, these findings highlight a complex regulatory network in which vessels, Notch, hypoxia, and Hippo/YAP1 shape endothelin receptor expression in glioma cells with the contrasting responses of IDH1-mutant LGG275 and IDH1-wild-type Gb7 underscoring glioma subtype–specific regulatory mechanisms.

### Bioinformatic and spatial analyses reveal a vascular-associated subpopulation of EDNRA⁺ tumor cells in GB

Because Notch signaling and hypoxia both upregulate EDNRA, we next investigated whether EDNRA⁺ tumor cell subpopulations exist in gliomas. Transcriptomic datasets consistently showed higher EDNRA expression in high-grade compared with low-grade gliomas (**Fig. S1A-B**), a trend we confirmed by IHC on a glioma tissue array [37] (**Fig. S10A**) and by IF on one grade II and one GB gliomas (**Fig. S10B**). In GB, EDNRA staining was stronger and enriched in vascular structures (**Fig. S10B**), co-localizing with CD31, consistent with its expression by mural cells such as pericytes and smooth muscle cells [65]. scRNA-seq analyses of GB further confirmed robust EDNRA expression in vascular cells (**Fig. S3A**) [46]. Unexpectedly, a small subset of tumor cells also expressed EDNRA in this dataset (**Fig. S3A, red arrow**), a finding independently reproduced in another GB dataset (**Fig. S3B, red arrow**) [13]. Given previous reports that some vascular-like cells in GB may derive from tumor cells [66], we hypothesized that these EDNRA⁺ tumoral cells could represent a vascular-like tumoral sub-population supporting the increased vascularization found in aggressive gliomas. This hypothesis is further supported by our earlier finding that Notch activation drives Gb7 cells toward a vascular-like phenotype and perivascular localization [45]. To test this possibility in patient tumors, we analyzed two other independent GB scRNA-seq datasets: (1) The study by Xi et al. shows that tumor cells co-purifying with vascular cells, express high EDNRA and low EDNRB (**Fig. S10C**) [67]; (2) Wang et al. (2023) identified a tumor subpopulation (Cluster 7) characterized by high EDNRA expression, enrichment for angiogenesis and EMT genes, and association with recurrent GBM, which they proposed to be linked to blood vessels [68]. To further assess this latter possibility, we mapped cells with this Cluster 7 transcriptomic profile using spatial transcriptomic data from three GB patients from Ravi et al, [69]. As expected, EDNRB expression was generally higher than EDNRA across tumor sections (**Fig. S11A**). However, spatial mapping of cells with Cluster 7 gene signature revealed that these EDNRA^+^ tumor cells co-localized with endothelial and pericyte clusters, primarily in perivascular regions (**Fig. S11B**). Collectively, these scRNA seq and spatial transcriptomic data establish the existence of a minor but distinct population of EDNRA⁺ tumor cells associated with the vasculature in GB.

## Discussion

In this study, we revisited endothelin signaling in diffuse gliomas using a newly established panel of serum-free cultured glioma cell lines representative of the major pathological subtypes— astrocytoma, oligodendroglioma, and glioblastoma. We combined this cellular resource with a broad range of high-throughput approaches, including bulk and single-cell RNA sequencing, spatial transcriptomics, proteomics, electrophysiology, and patient-derived tumor extracts. A synthesis of these results is presented in Figure 7.

**Figure 7.**
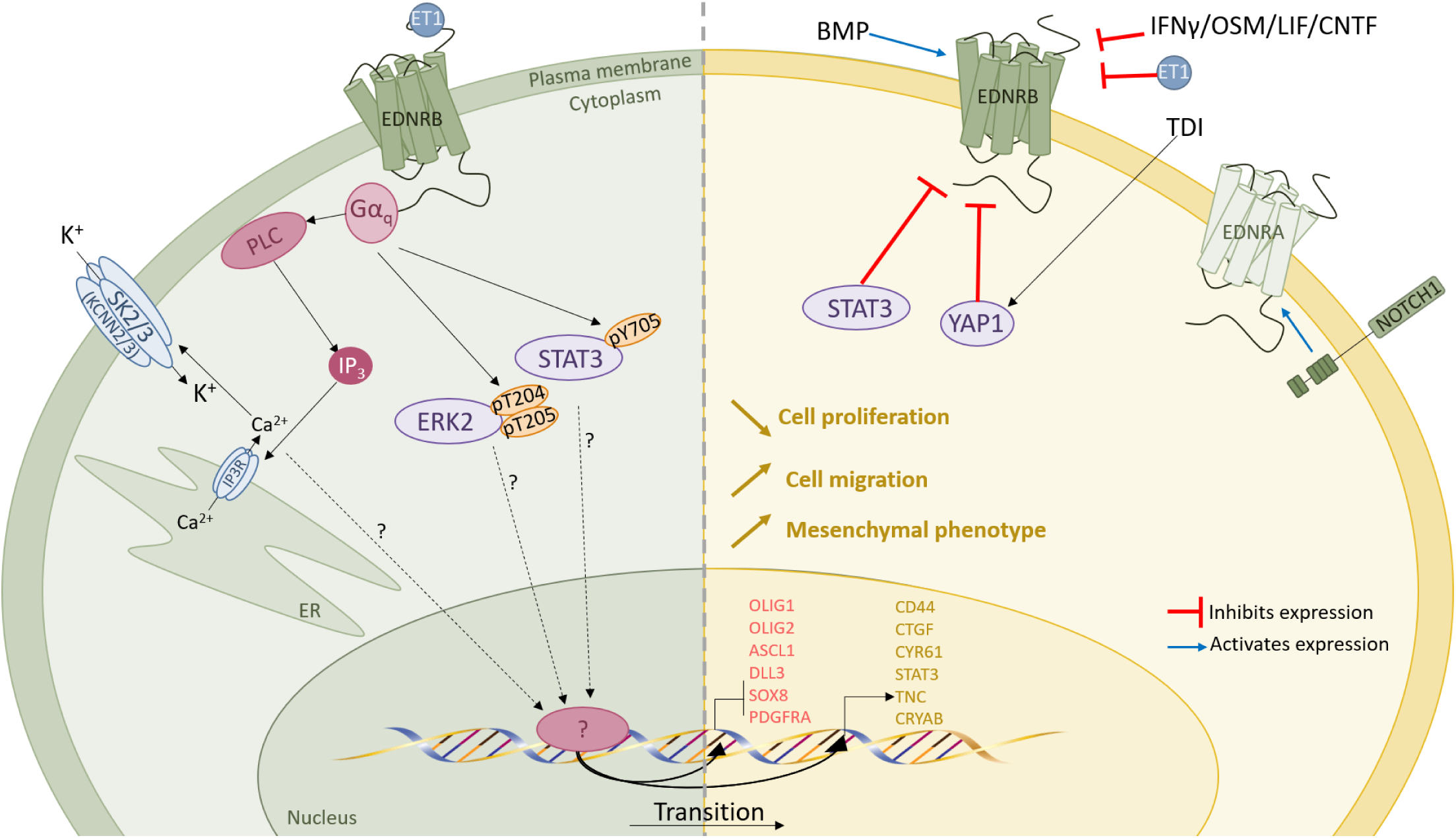
Proposed model of endothelin signaling and regulation in diffuse glioma cells. EDNRB is predominant over EDNRA, and its activation by ET-1 stimulates downstream pathways including Ca²⁺ influx, SK K⁺ channels, ERK, and STAT3, thereby influencing glioma cell proliferation, migration, and the acquisition of a mesenchymal phenotype. BMP signaling upregulates EDNRB, whereas STAT3, IFNs and Hippo/YAP1 repress its expression. In glioblastoma, Notch1 activation upregulates EDNRA.

Our data shows that EDNRB is the dominant endothelin receptor in glioma cell lines and patient tumors. It is enriched in tumor cells with an astrocyte-like phenotype, and its expression increases following growth-factor withdrawal or exposure to BMPs. By contrast, EDNRB is repressed by interferons, IL-6-family cytokines, endothelins themselves, and activation of the Hippo–YAP pathway. Taken together, these findings indicate that EDNRB is differentially regulated by multiple, mechanistically distinct signaling cues present in the glioma microenvironment, which likely contributes to the coexistence of EDNRB-positive and EDNRB-negative tumor cell populations within gliomas.

EDNRA, in contrast, is primarily expressed by vascular cells, as shown in previous studies [22], [68]. However, our bioinformatic analyses revealed that a small subpopulation of GB cells also express EDNRA. Actually, in the GB cell line we examined (Gb7), EDNRA expression was inducible by Notch1 signaling and, to a lesser extent, by hypoxia. EDNRA expression in Gb7 cells could also be detected *in vivo* by immunoPET imaging in a preclinical orthotopic xenograft model [70], [71]. This demonstrates the capacity of GB cells to upregulate EDNRA in response to specific pathways and environment. This capacity was absent in the IDH1-mutant LGG275 line, where we suspect that the DNA and chromatin hypermethylation characteristic of the IDH1-mutant context precludes EDNRA activation.

Functionally, activation of EDNRB by endothelins reduces proliferation, enhances migration, and reprograms gene expression toward a more mesenchymal profile. This shift coincides with (i) repression of OPC-specific genes such as OLIG1 and (ii) activation of several downstream pathways: Ca²⁺ signaling, ERK, STAT3, and Ca²⁺-gated K⁺ channels. The strong induction of STAT3 is particularly noteworthy, given its established role in driving mesenchymal transition in glioma [16], [17]. Our antiproliferative findings contrast with earlier reports that described endothelins as mitogenic in gliomas [72]. A likely explanation is methodological: most of those studies relied on serum-cultured cell lines, where undefined serum components can profoundly reshape signaling networks and may mask the growth-inhibitory effect of EDNRB activation observed under serum-free, defined conditions.

In conclusion, G protein-coupled receptors regulate many facets of cell behavior, yet their involvement in the proneural-to-mesenchymal transition in glioma remains poorly documented [73]. To date, CXCR4-the GPCR for SDF-1/CXCL12-is one of the few receptors shown to drive PMT in this tumor type. Our data now add EDNRB to this list, revealing a new GPCR-mediated mechanism that governs glioma cell plasticity. Glioma cells situated in close proximity to blood vessels exhibit lower proliferative activity [21]. Since vascular endothelial cells are the principal source of endothelins in the brain, it is plausible that vessel-derived endothelins help enforce this growth restraint. Interestingly, Kaplan–Meier analyses indicate that patients with high EDNRB expression exhibit longer survival, in contrast to EDNRA (**Fig. S1E**). This suggests that loss of EDNRB may facilitate glioma cells in evading anti-proliferative signals from the tumor microenvironment. A limitation of our study is that we were unable to directly test this hypothesis in vivo.

---

**Figure S1.**
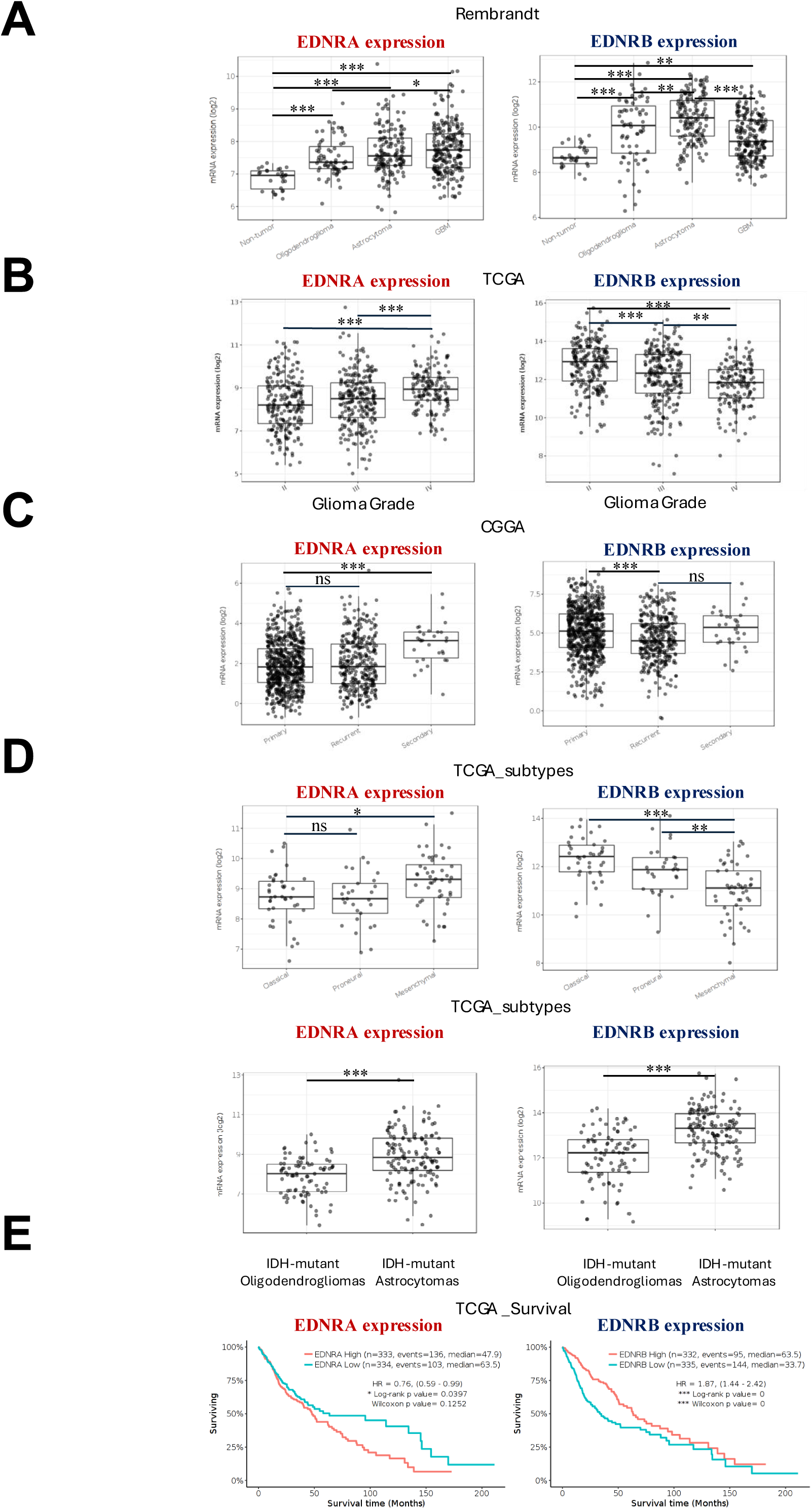
Endothelin receptor gene expression across human glioma datasets (related to Fig. 1). mRNA levels (log₂) for *EDNRA* and *EDNRB* were obtained from the GlioVis portal (accessed 3 Jan 2025) and plotted as box-and-whisker plots; each dot represents one tumor sample. Statistical significance was calculated in GlioVis using one-way ANOVA with Tukey’s post-hoc test; * p <0.05, ** p <0.01, *** p <0.001, ns = not significant. **A**. Rembrandt microarray cohort: non-tumoral brain (n=28) vs oligodendroglioma (ODG, n=67), astrocytoma (AST, n=147), and glioblastoma (GB, n=219). **B**. TCGA pan-glioma RNA-seq cohort: stratified by WHO tumor grade II (n=226), III (n=244), and IV (n=150). **C**. CGGA RNA-seq cohort: primary (n=651) vs recurrent (n=333) gliomas. **D**. TCGA molecular subtypes: *(top)* IDH-wild-type GB—Classical (n=199), Mesenchymal (n=166), Proneural (n=163); *(bottom)* IDH-mutant lower-grade gliomas—oligodendroglioma (n=85) vs astrocytoma (n=141). **E.** TCGA pan-glioma Kaplan-Meier estimator survival analysis of EDNRA and EDNRB expression (high and low) over 200 months (Cutoff: median). Statistical significance was calculated in GlioVis and is displayed on the graph.

**Figure S2.**
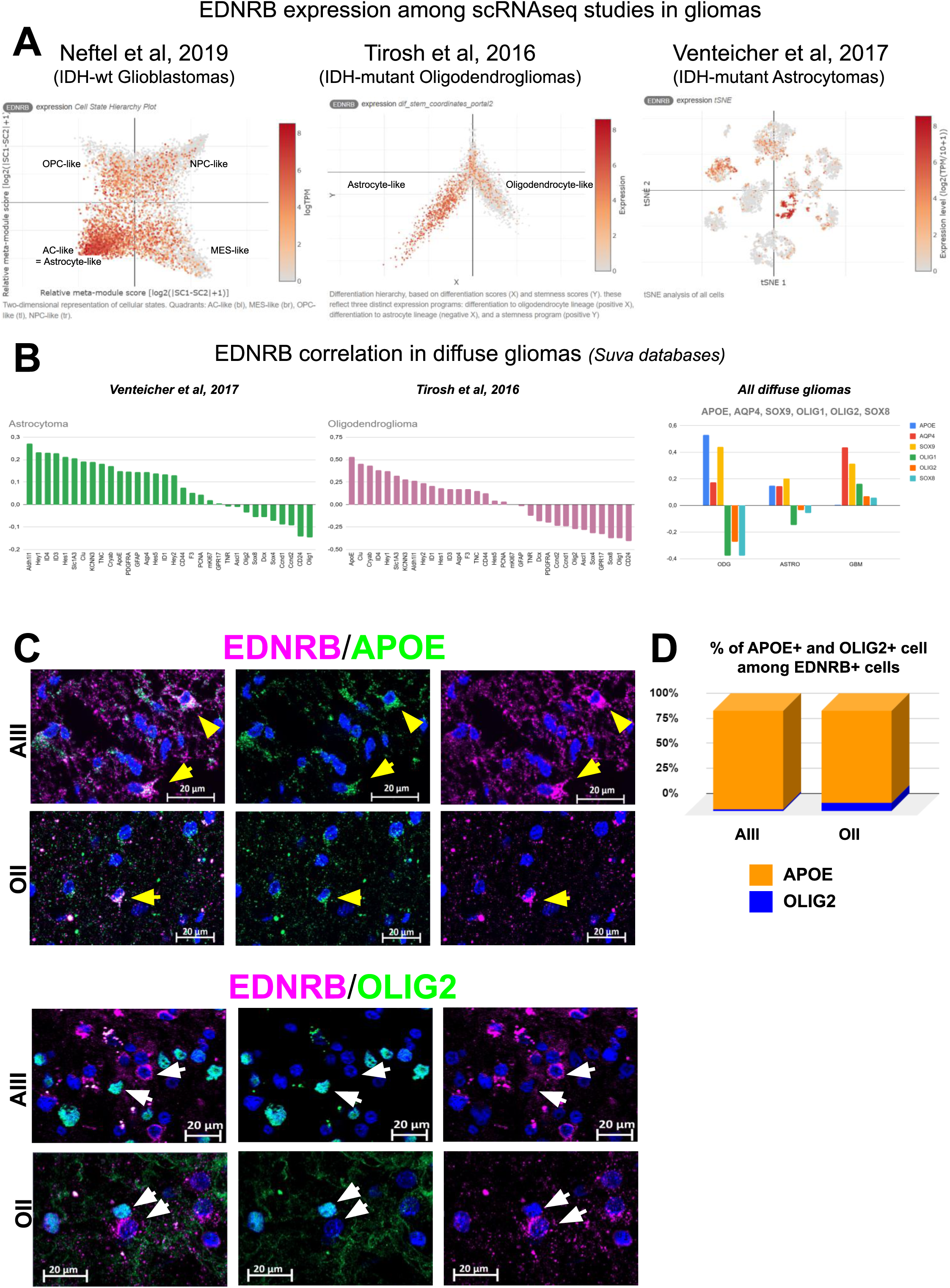
EDNRB is preferentially expressed in astrocyte-like glioma cells (related to Fig. 2). **A.** Single-cell RNA-seq from three studies showing EDNRB expression by glioma cell subtype. (*Left*) Neftel et al., 2019 (IDH–wild-type GB): enrichment in astrocyte-like (AC-like) cells. (*Middle*) Tirosh et al., 2016 (IDH–mutant oligodendrogliomas): higher in AC-like than oligodendrocyte-like cells. (*Right*) Venteicher et al., 2017 (IDH–mutant astrocytomas): heterogeneous expression across clusters. Color scale: red = high, white = low. **B.** (*Left*) Correlation between EDNRB expression and lineage-associated genes in diffuse IDH1-mutant astrocytomas and oligodendrogliomas, based on scRNA-seq data from Venteicher et al., 2017[9]. Correlation coefficients were manually retrieved from the study’s online database. In astrocytomas, EDNRB shows positive correlations with astrocytic markers and negative correlations with oligodendrocytic genes, indicating an astrocyte-like profile—similar to that observed in GB. (*Right*) Right: correlation of EDNRB with APOE, SOX9, AQP4, OLIG1, OLIG2, SOX8 in ODG, ASTRO, and GB, from Suvà group datasets ([9], [10], [13]). **C**. Representative immunofluorescence of astrocytomas (AIII) and oligodendrogliomas (OII) stained for EDNRB (*purple*), APOE or OLIG2 (*green*), nuclei (Hoechst, *blue*). *Yellow* arrowheads: EDNRB⁺/APOE⁺ cells; *white* arrows: EDNRB⁻/OLIG2⁺ or EDNRB⁺/OLIG2⁻ cells. Scale bars: 20 µm. **D**. Quantification of APOE⁺ and OLIG2⁺ cells among EDNRB⁺ cells in AIII and OII shows most EDNRB⁺ cells are APOE⁺, not OLIG2⁺.

**Figure S3.**
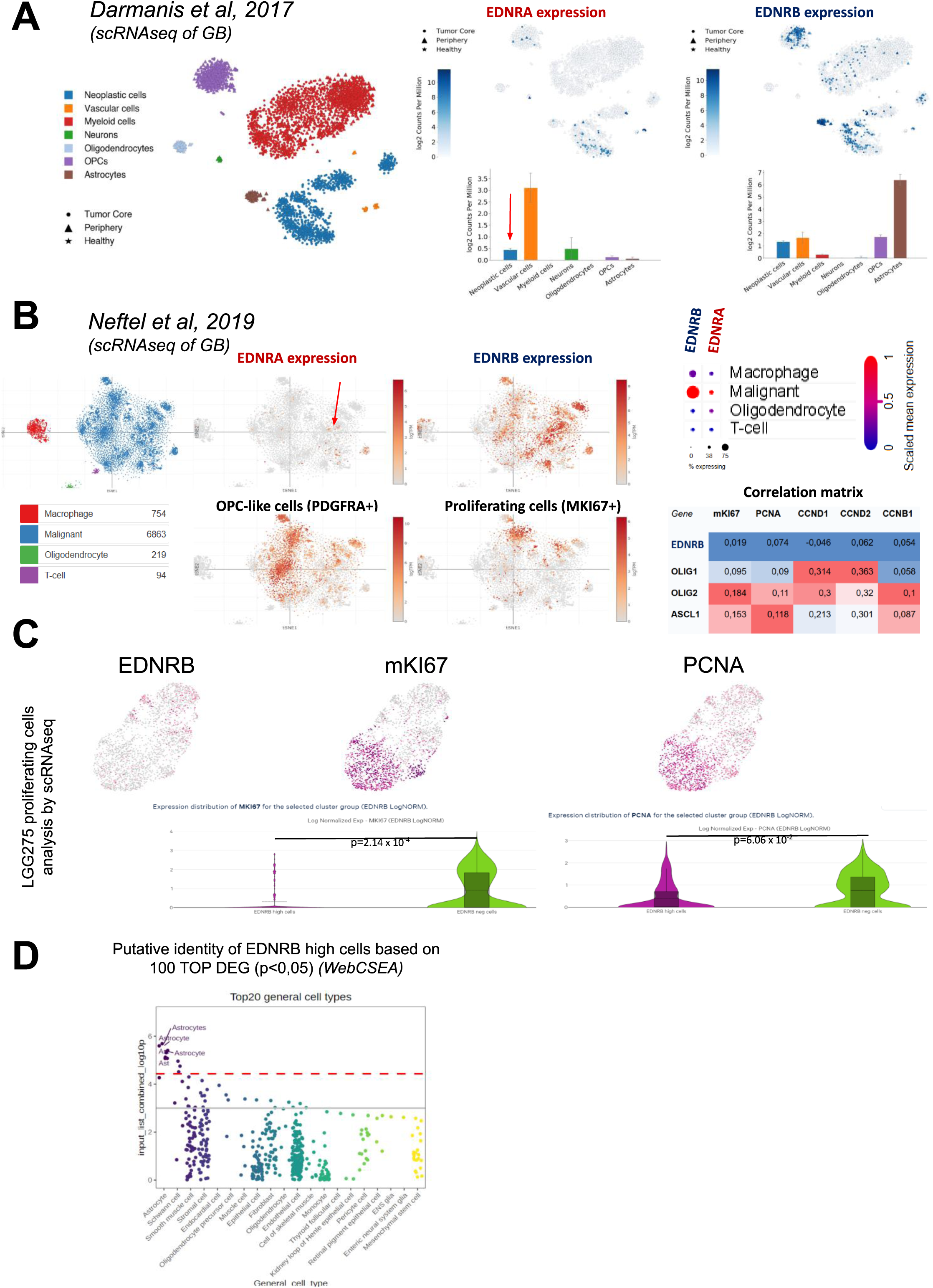
EDNRB expression defines a low-proliferative, astrocyte-like cell population in gliomas. **A.** scRNA-seq analysis of glioblastoma (Darmanis et al., 2017) showing cell-type composition and EDNRA/EDNRB expression patterns.[46] (*Left*) UMAP plots display major cell populations (color-coded) from tumor core, periphery, and healthy tissue. (*Right*) EDNRA expression is predominantly enriched in vascular/mural cells, with a small subpopulation of tumor cells also expressing EDNRA (red arrow). EDNRB shows highest expression in astrocytes and is also present in tumor and vascular cells. Bar graphs indicate log₂ counts per million (mean ± SEM) for each cell type. **B.** scRNA-seq analysis of glioblastoma (Neftel et al., 2019). (*Left)* t-SNE plot showing major cell types (color-coded). (*Middle*) Single-cell expression maps of EDNRA, EDNRB, PDGFRA (OPC marker), and MKI67 (proliferation marker). Tumor cells preferentially express EDNRB, but a small population also expresses EDNRA (red arrow). (*Top right*) Dot plot summarizing EDNRA and EDNRB expression levels and percentages across the four main cell types in GB. (*Bottom right*) Pearson correlation matrix between lineage-associated genes (EDNRB, OLIG1, OLIG2, ASCL1) and proliferation markers (MKI67, PCNA, CCND1, CCND2, CCNB1), showing minimal association of EDNRB with proliferation compared to oligodendrocyte-lineage genes. **C.** scRNA-seq analysis of LGG275 cells. UMAPs showing EDNRB, MKI67, and PCNA expression (*pink* = high, *grey* = low). Violin plots compare proliferation marker levels between EDNRB-high and EDNRB-negative cells. Tests: Benjamini–Hochberg adjusted p-values. EDNRB-high cells show reduced proliferation. **D.** WebCSEA (Web-based Cell-type-Specific Enrichment Analysis, UTHealth) cell-type enrichment of EDNRB-high LGG275 cells. Top 100 differentially expressed genes (p <0.05) were analyzed, Top 20 general cell types displayed on the graph are revealing significant enrichment for astrocytes (*purple*; *red* dashed line = adjusted p <0.01).

**Figure S4.**
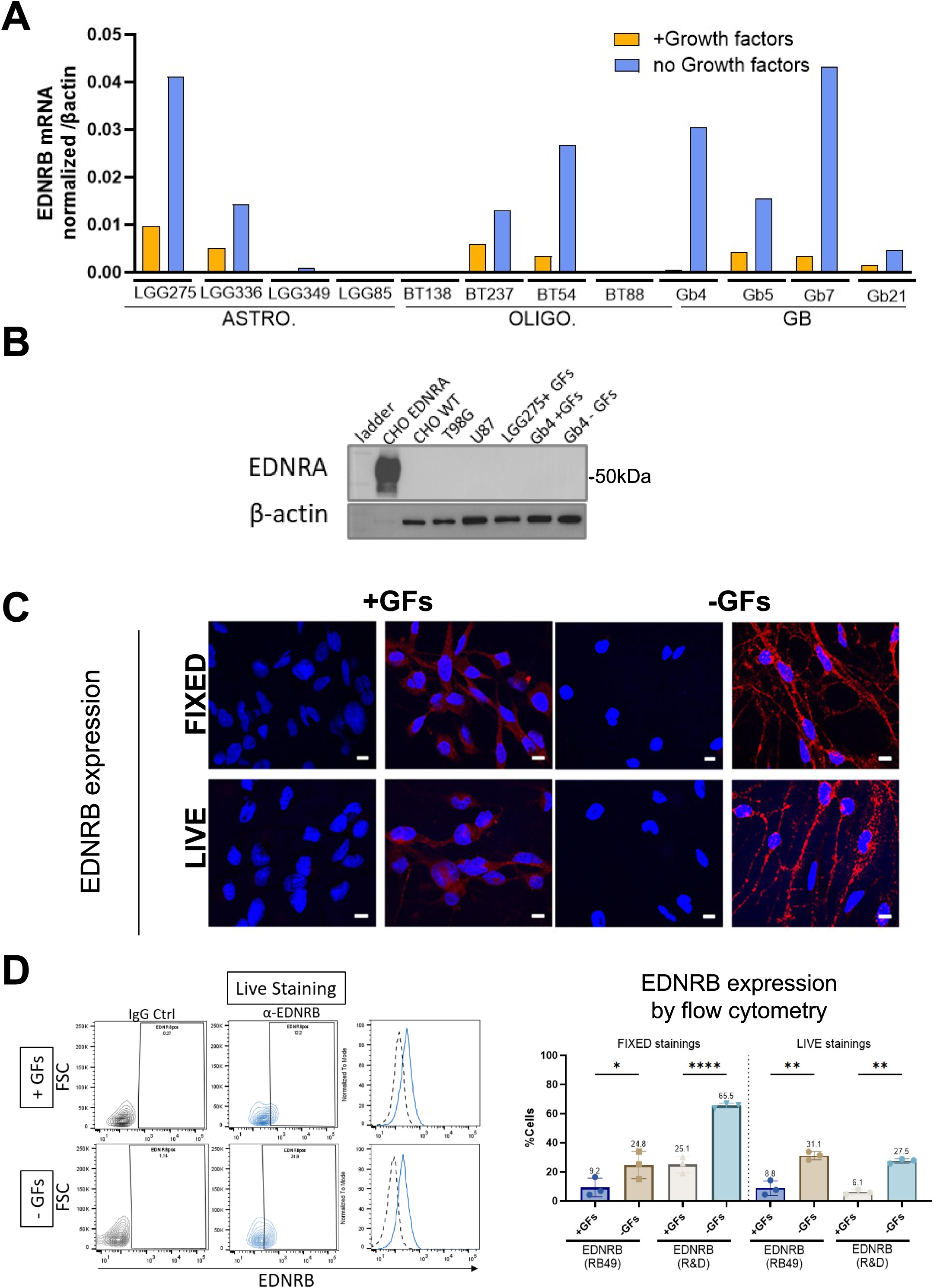
EDNRB is the predominant endothelin receptor and is expressed at the cell surface of diffuse glioma cultures (related to Fig. 2). **A**. RT–qPCR validation of EDNRB expression (confirming RNA-seq from Fig. 2A) in 11 patient-derived glioma lines and one additional GB line (Gb5) [74], cultured with (+GFs) or without (–GFs) growth factors. **B**. Western blot of EDNRA in glioma lines (T98G, U87, LGG275, Gb4) and CHO cells. EDNRA was detected only in EDNRA-overexpressing CHO cells (positive control) and was absent from glioma lines, consistent with Fig. 2A. **C**. Immunofluorescence of LGG275 cells stained for EDNRB (red; RB49 antibody targeting an extracellular epitope [75]) under fixed or live conditions. Nuclei were counterstained with Hoechst (*blue*). EDNRB localizes to the cell surface, with stronger labeling in –GF conditions. Scale bar, 10 µm. **D.** Flow cytometry of EDNRB surface expression in fixed and live LGG275 cells cultured ±GFs, using RB49 and a commercial antibody (R&D Systems, Cat. #AF4496). Bar graphs show mean ± SD of % EDNRB⁺ cells (n = 3). Statistics: one-way ANOVA with Tukey’s test; *p <0.05, **p <0.01, ****p <0.0001.

**Figure S5.**
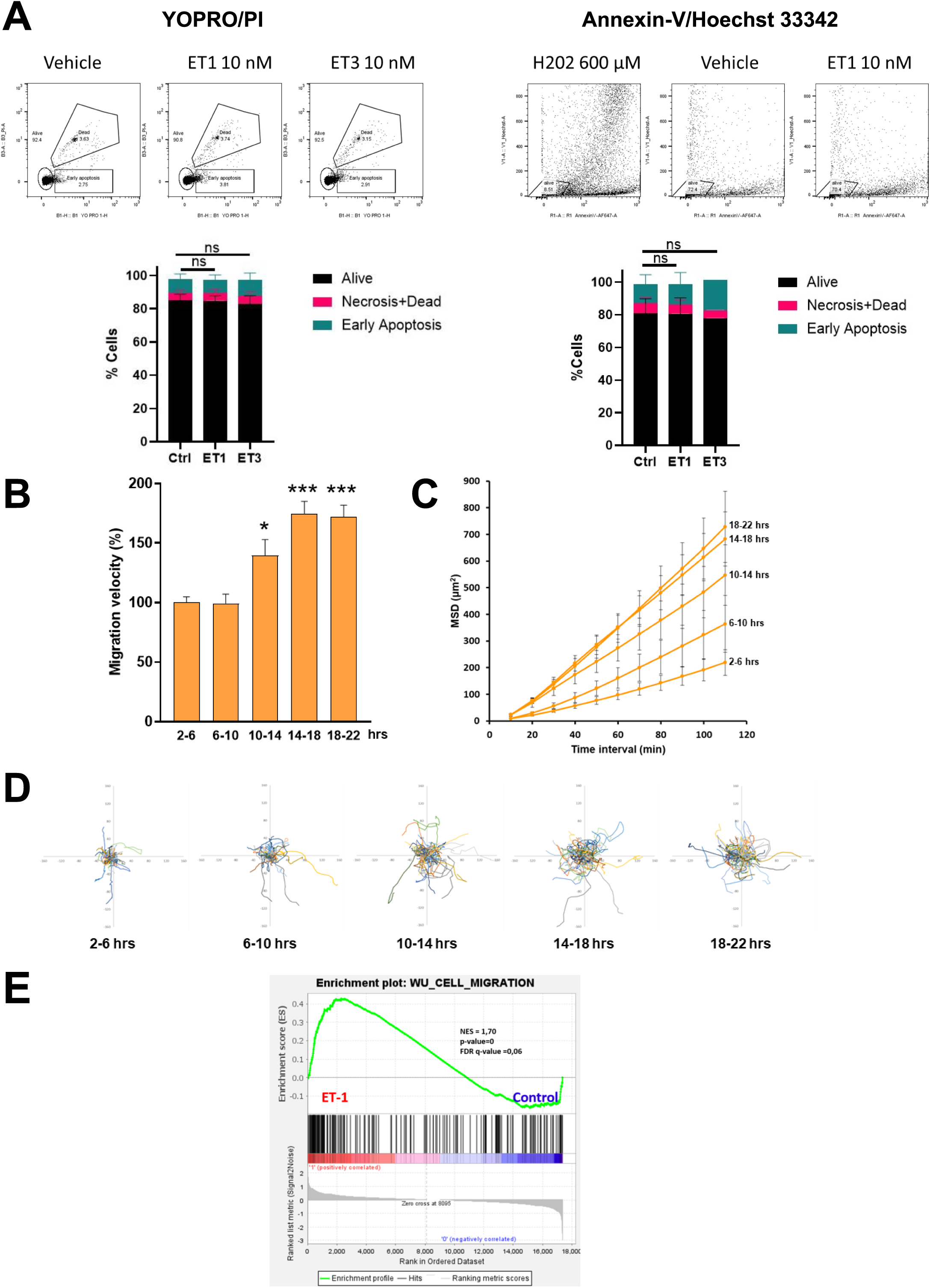
Further characterization of ET-1 effects on migration and cell death (related to Fig. 3) **A.** Flow cytometry analysis of cell death in LGG275 cells treated with ET-1 or ET-3. Cell death was assessed using (left) YO-PRO/Propidium Iodide or (right) Annexin-V/Hoechst 33342 staining. H₂O₂ (600 µM) served as a positive control (not shown). Data are mean ± SD (n = 3 independent experiments for ET-1 in both assays; n = 3 independent experiments for ET-3 with YO-PRO/PI; n = 1 for ET-3 with Annexin-V/Hoechst). ns, not significant (ANOVA with Tukey’s post-hoc test). **B.** Time-lapse analysis of LGG275 migration after ET-1 treatment (10 nM). Migration velocity was tracked over 22 h, showing a significant increase from 10 h onward (*p <0.05 to ***p <0.001). **C.** Mean square displacement (MSD) of LGG275 cells measured over successive 4 h intervals (2–22 h) after ET-1 treatment, showing increased displacement and migration efficiency over time. **D.** Migration trajectories of individual LGG275 cells in consecutive 4 h intervals following ET-1 treatment, showing a progressive displacement from the point of origin over time **E.** GSEA of RNA-seq data from ET-1–treated LGG275 cells showing significant enrichment of the *WU_Cell_Migration* gene set. NES, nominal *p*-value, and FDR *q*-value are indicated.

**Figure S6.**
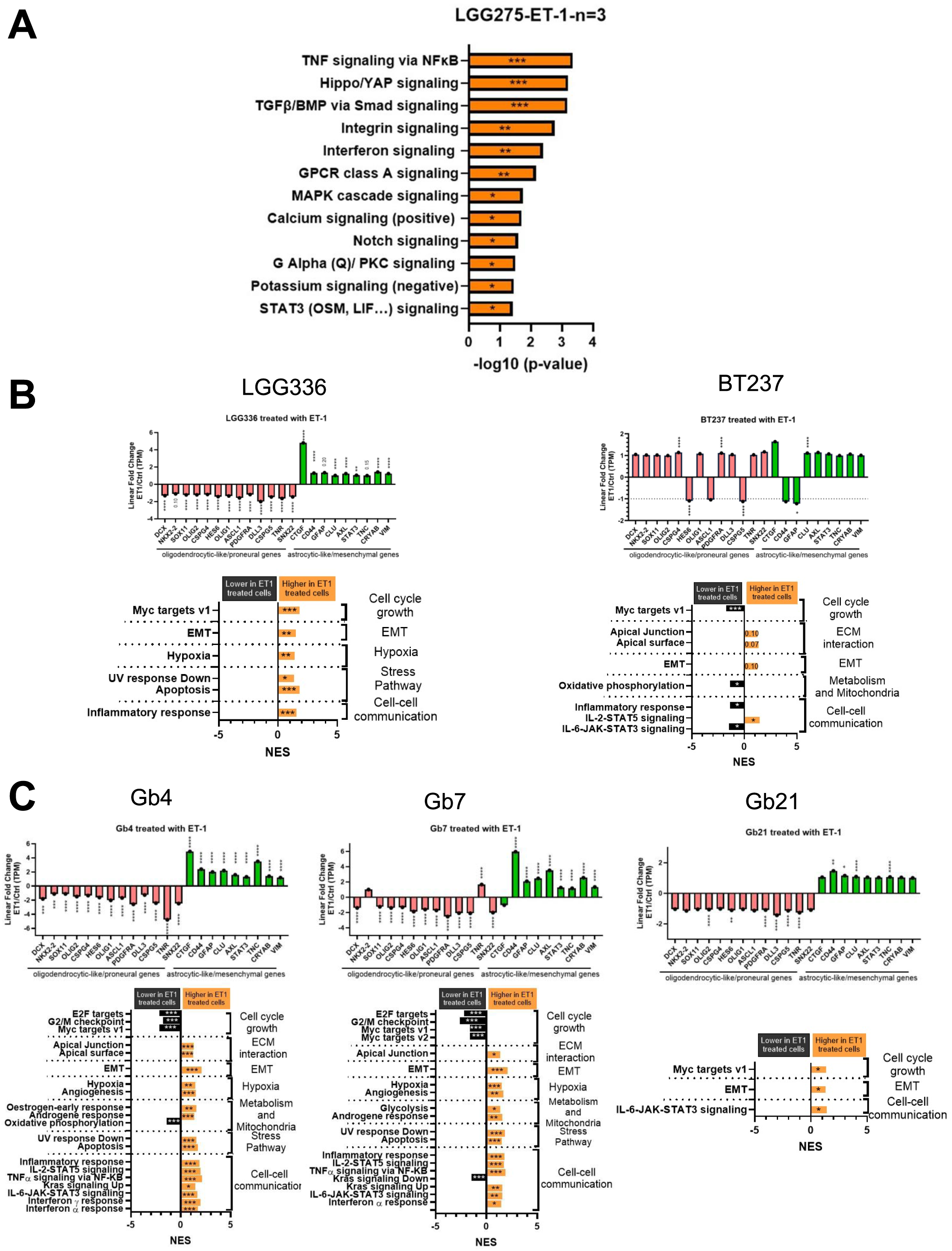
RNA-seq analysis of glioma cell lines treated with ET-1 (related to Fig. 4). **A.** Pathway enrichment (EnrichR) from 65 DEGs in LGG275 cells after ET-1 treatment (n = 3 independent experiments)[55]. Bars show –log₁₀(p) values; significance: p <0.05 (*), <0.01 (**), <0.001 (***). **B–C.** Analyses of Bulk RNA-seq of ET-1–treated glioma lines: *(B)* IDH1-mutant (LGG336, BT237) and *(C)* IDH–wildtype GB (Gb4, Gb7, Gb21) (n=1 per condition). *Top panels*: Fold-change of lineage-associated genes (oligodendrocytic/proneural = purple; astrocytic/mesenchymal-like = green). *Bottom panels*: GSEA for each line; all processes FDR q <0.25. Significance: p <0.05 (*), <0.01 (**), <0.001 (***). ET-1 promotes astrocytic/mesenchymal transcriptional reprogramming in both IDH1-mutant and IDH–wildtype glioma cells.

**Figure S7.**
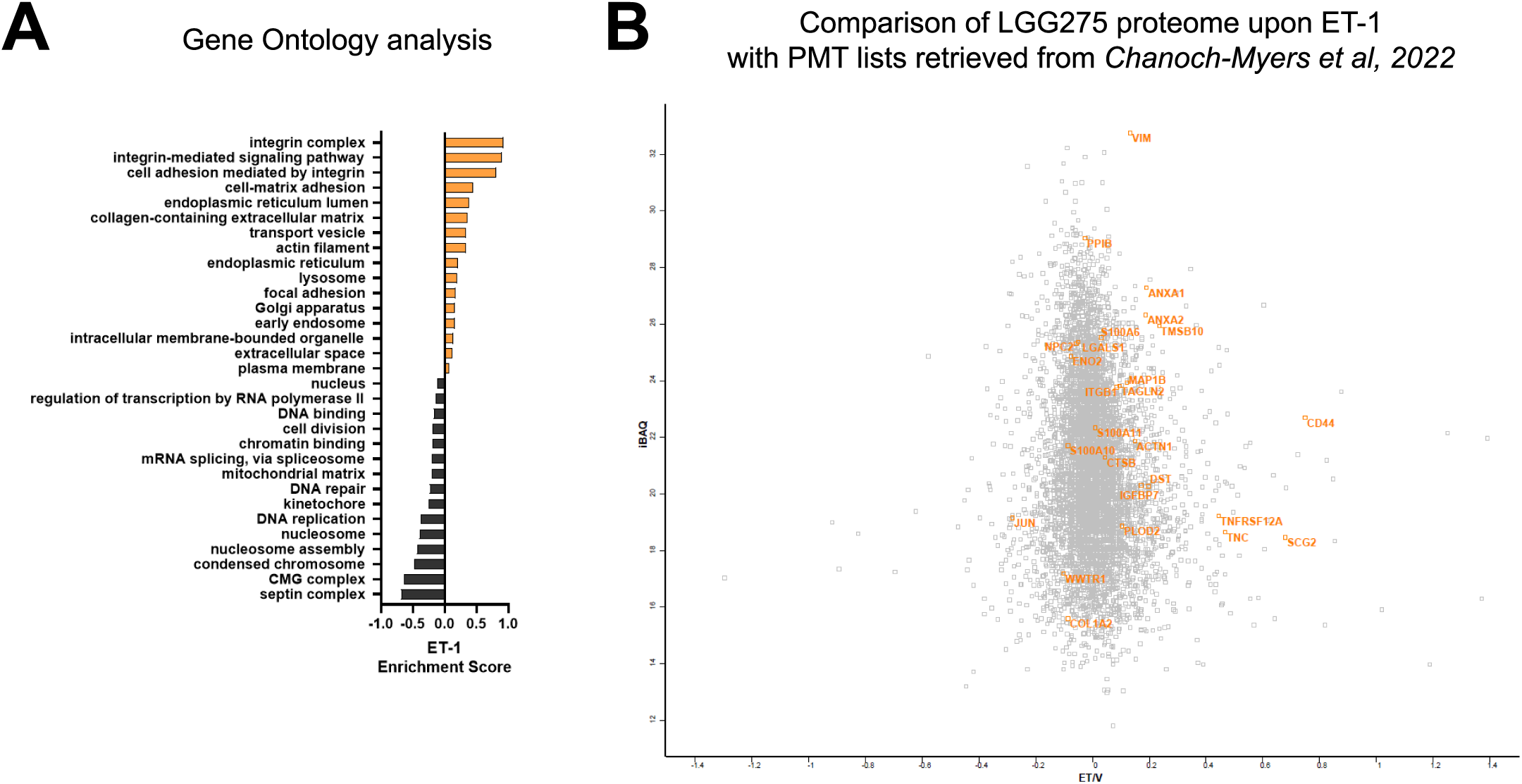
ET-1 induces shifts in the proteomic profile associated with mesenchymal transition signatures in LGG275 cells (related to Fig. 4). **A**. Gene Ontology (GO) term enrichment analysis of proteomic profiles from LGG275 cells treated with ET-1 versus vehicle, based on five independent experiments and analyzed by 1D annotation enrichment with Fisher’s exact test. Orange bars indicate positive enrichment, black bars negative enrichment. **B**. Scatter plot of proteins quantified by TMT-based mass spectrometry (ET-1 vs vehicle) (same as Fig 4H). The x-axis shows the log₂ ratio (ET/V) and the y-axis the iBAQ value. Proteins in orange correspond to 26 of the 28 proneural-to-mesenchymal transition (PMT) markers defined by Chanoch-Myers et al., 2022 [14]; 73% (19 out of 26) of these are upregulated following ET-1 treatment, consistent with a mesenchymal-like shift.

**Figure S8.**
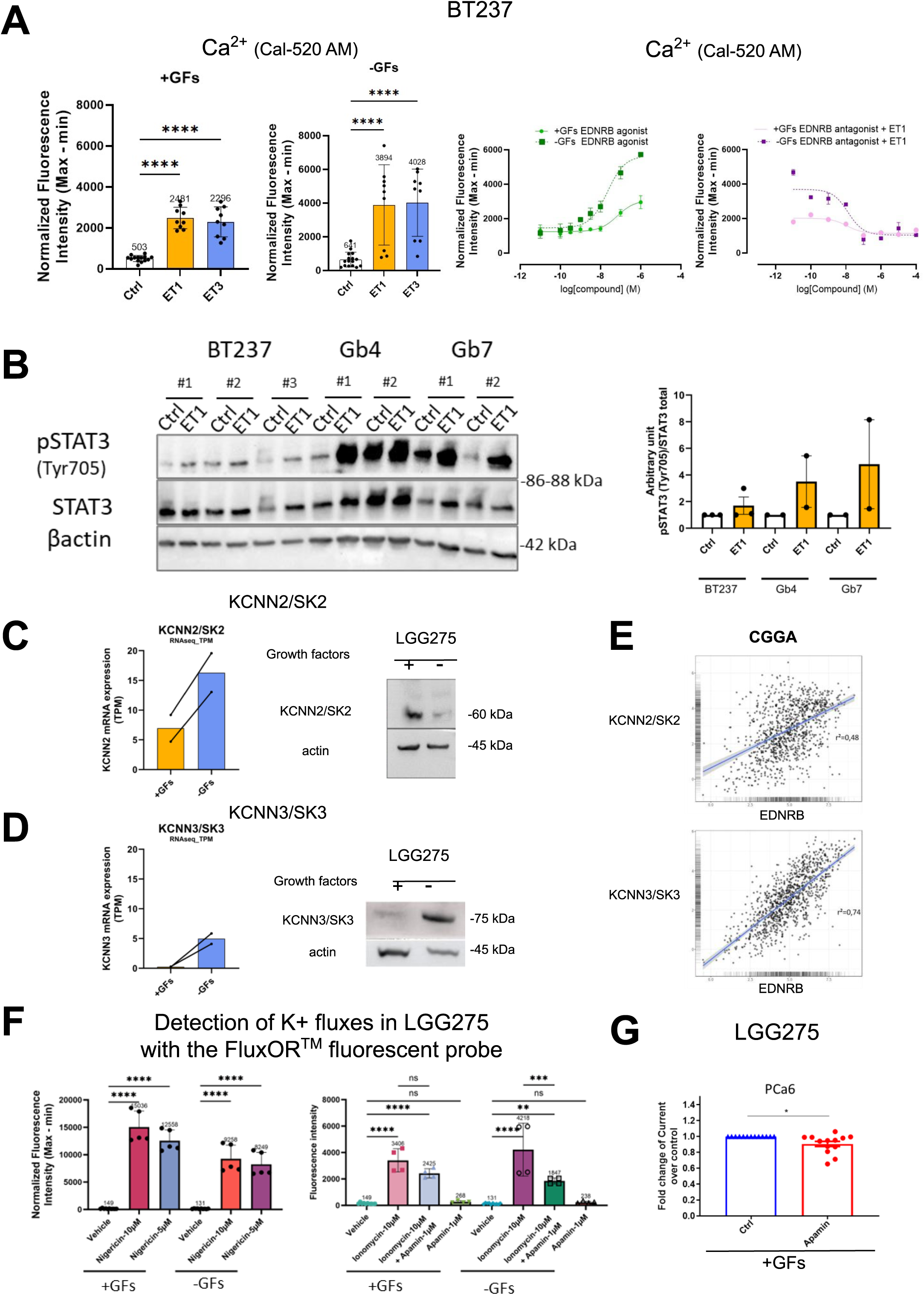
Signaling stimulated by endothelins in glioma cell lines (related to Fig. 5) **A**. ET-1– induced Ca²⁺ responses in BT237 IDH1-mutant oligodendroglioma cells (related to Fig. 5 C). *Left*: Intracellular Ca²⁺ fluxes (Cal-520 AM) in control, ET-1, or ET-3–treated cells cultured with (+GFs) or without (–GFs) growth factors. Data: mean ± SD, n = 3 independent experiments; one-way ANOVA/Tukey; ****p <0.0001. *Right*: Dose–response curves for the EDNRB agonist IRL-1620 and the EDNRB antagonist BQ-788 plus EC₈₀ ET-1 (4 nM) with or without GFs. Curves: non-linear regression; one representative of three experiments shown. **B.** ET-1–stimulated STAT3 signaling in glioma cell lines. *Left*: Western blot of phosphorylated STAT3 (pSTAT3, Tyr705), total STAT3, and β-actin in BT237 (n = 3), Gb4 (n = 2), and Gb7 (n = 2) cells treated with ET-1 or vehicle control. *Right*: Quantification of pSTAT3 normalized to total STAT3; values are expressed as arbitrary units (A.U.) relative to control (set to 1). **C–D.** Expression of KCNN2/SK2 and KCNN3/SK3 in LGG275 cells. RNA (TPM, RNA-seq; n = 2) and protein (Western blot) levels in cells cultured with (+GFs) or without (– GFs) growth factors. Actin was used as a loading control. **E.** Correlation of EDNRB expression with KCNN2 and KCNN3 in gliomas. Scatter plots from the CGGA dataset (n = 1,016) showing Pearson’s correlation between EDNRB and KCNN2 (top) or KCNN3 (bottom) expression. r² and p-values =0, (two-sided) from GlioVis database. **F.** Validation of FluxOR™ probe for detecting K⁺ fluxes in LGG275 cells (related to Fig. 5 F). *Left*: Normalized fluorescence intensity (Max–Min) after treatment with the K⁺ ionophore nigericin (10 µM or 5 µM) under +GFs or –GFs conditions. *Right*: K⁺ flux induced by the Ca²⁺ ionophore ionomycin (10 µM) with or without the SK-channel blocker apamin (1 µM), under +GFs or –GFs conditions. Data: mean ± SD, n = 2 independent experiments; one-way ANOVA/Tukey. ****p <0.0001; ***p <0.001; **p <0.01; ns: not significant. **G.** Apamin-sensitive K⁺ currents in LGG275 cells (+GFs). Fold change in membrane current amplitude (normalized to control) recorded under whole-cell voltage-clamp with intracellular free Ca²⁺ buffered at ∼1 µM (PCa6) to activate SK channels. Spontaneous Ca²⁺-dependent K⁺ currents were significantly reduced by apamin (100 nM), a selective SK channel blocker. Mean ± SD (n =13 cells). Unpaired t-test; *p <0.05.

**Figure S9.**
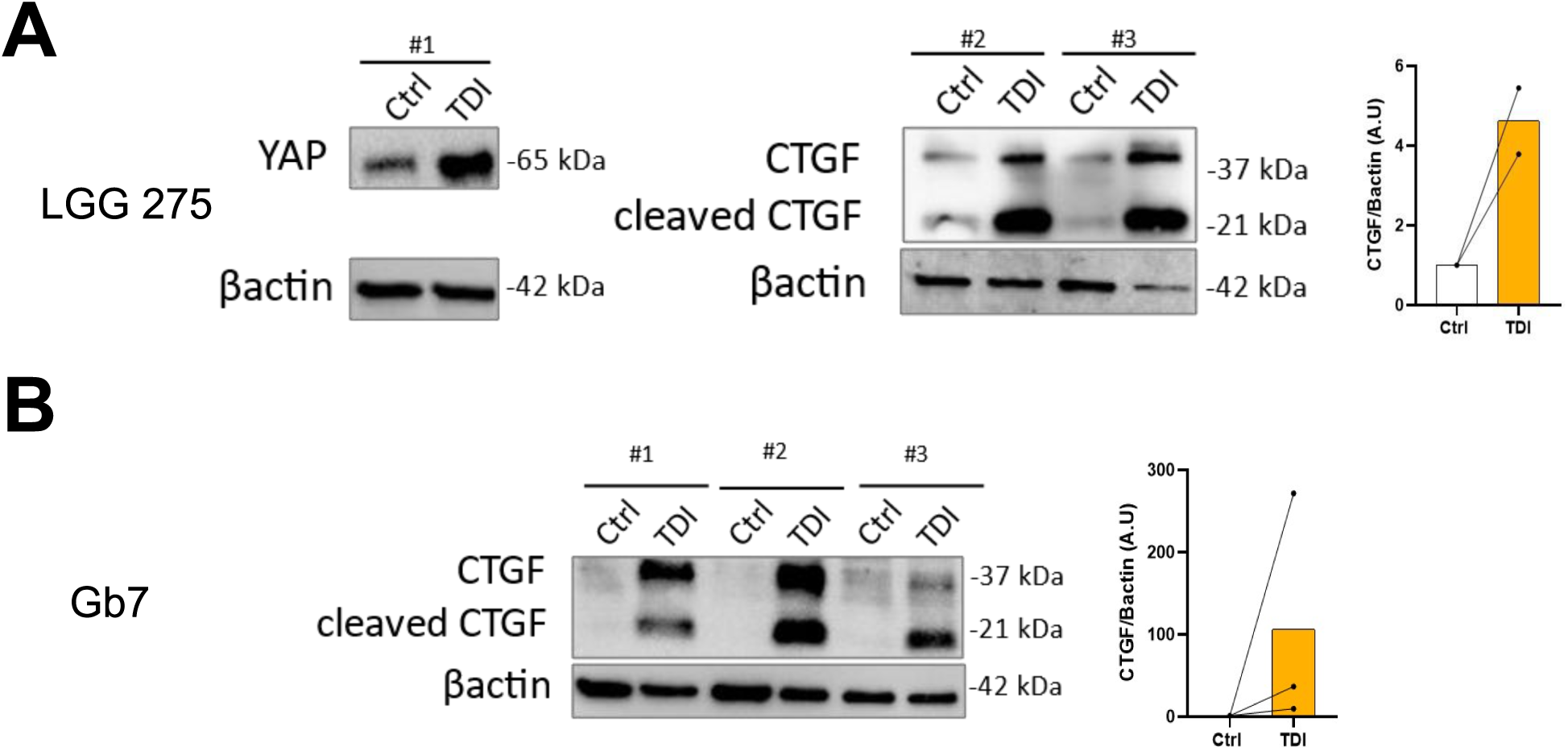
TDI activates the YAP pathway in diffuse glioma cells (related to Fig. 6). Western blot analysis of YAP and CTGF (full-length ∼37 kDa and cleaved ∼21 kDa) in LGG275 (A) and Gb7 (B) cells treated with TDI (1 µM) or vehicle (Ctrl). β-actin served as a loading control. Right panels: densitometric quantification of CTGF normalized to β-actin, expressed as arbitrary units (A.U.) with Ctrl set to 1. Data from 2–3 independent experiments.

**Figure S10.**
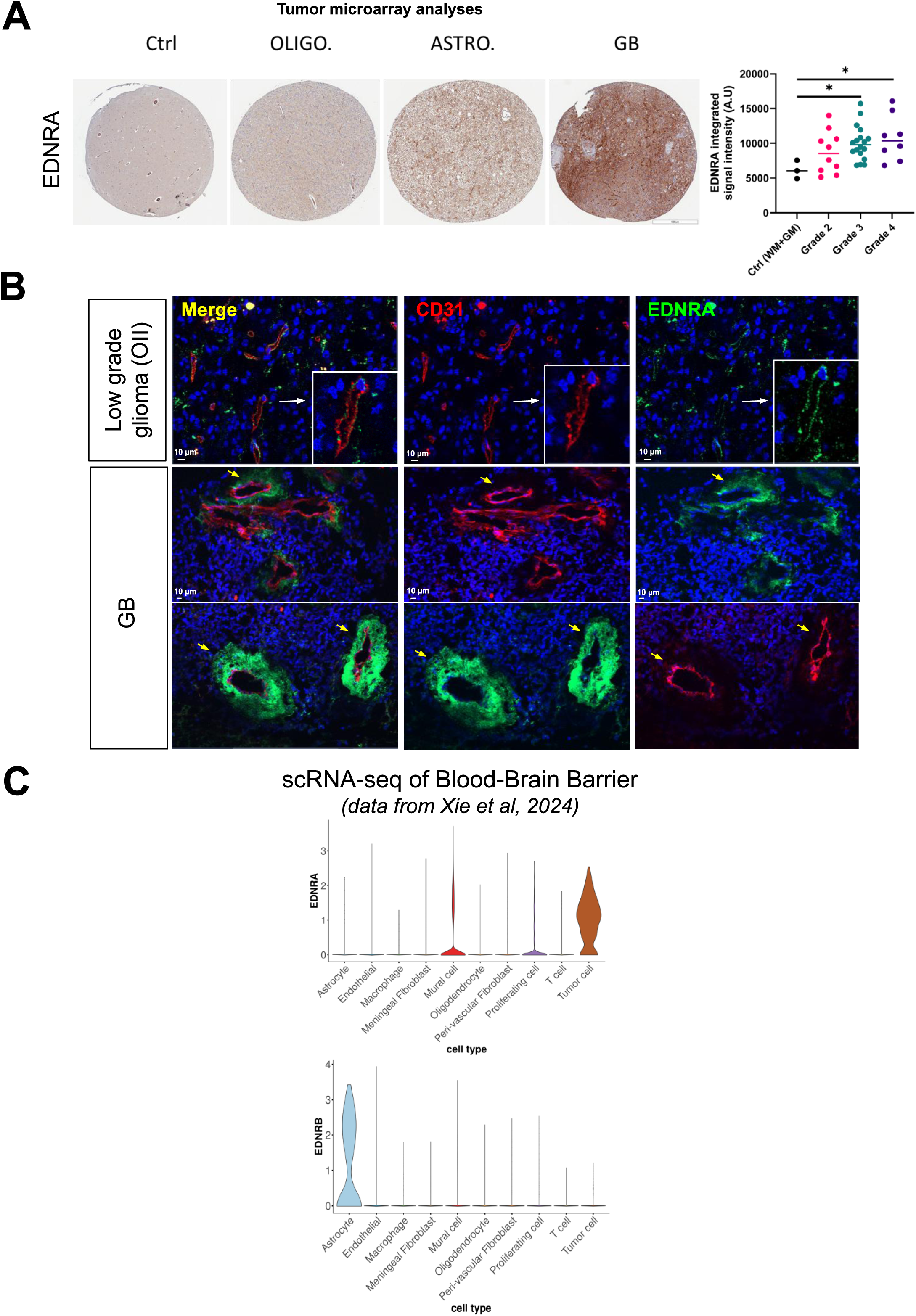
EDNRA expression increases with glioma grade. **A.** *(Left)* Immunohistochemistry on tumor microarrays (TMA [38]).) showing EDNRA in control brain (white + grey matter) and gliomas (oligodendrogliomas, astrocytomas, glioblastomas). Scale bar = 600 µm. *(Right)* Quantification of EDNRA integrated signal intensity (A.U.) in control and tumor samples (grades 2–4) from 39 patients. Data were analyzed by one-way ANOVA with Tukey’s post-hoc test; *p <0.05. **B.** Representative immunofluorescence images of oligodendroglioma (OII) and glioblastoma (GB) stained for EDNRA (*green*), CD31 (*red*), and nuclei (Hoechst, *blue*). Merged images (*left*) show EDNRA localization near CD31⁺ endothelial cells, with insets highlighting colocalization (*white* arrows, OII; *yellow* arrows, GB). EDNRA staining is stronger and more widespread around vessels in GB. Scale bars = 10 µm. **C.** Single-cell expression of EDNRA and EDNRB in vascular and tumor-associated cells. Violin plots showing normalized EDNRA (*top*) and EDNRB (*bottom*) expression across major brain cell types from purified vascular fractions of glioma tissues (scRNA-seq; Xie et al., 2024) [67]. These plots were adapted from the interactive resource available at https://scatlas.shinyapps.io/human_bbb_and_btb/. EDNRA is enriched in mural and vascular-associated tumor cells, whereas EDNRB is mainly detected in astrocytes. The absence of EDNRB in tumor cells here likely reflects the isolation strategy (MACS with CD31⁺ and PDGFRβ⁺ selection), which enriches for vascular/perivascular populations while excluding most tumor cells.

**Figure S11.**
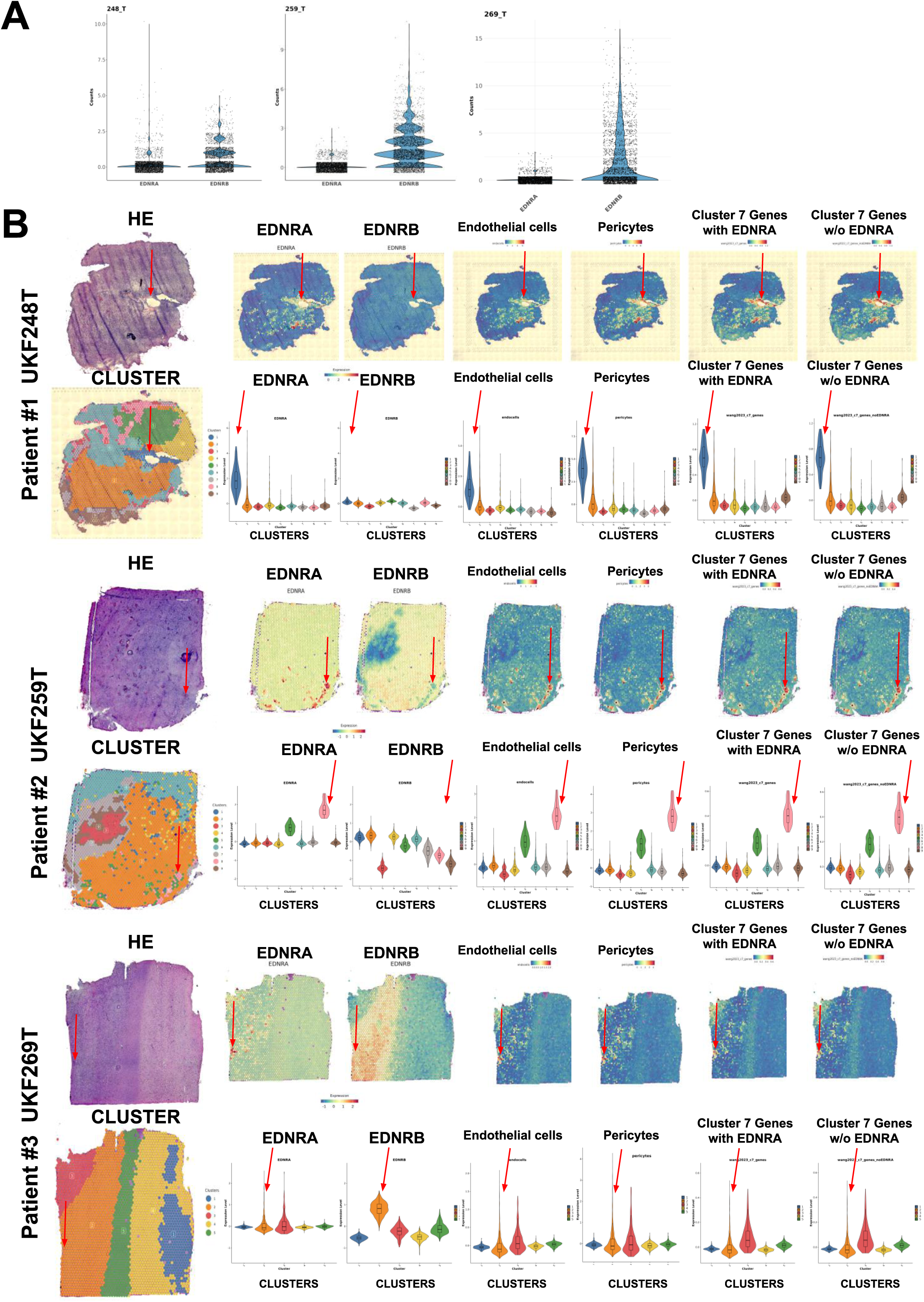
Spatial mapping of endothelin receptors in glioblastoma. **A**. Violin plots showing normalized transcript counts for EDNRA and EDNRB in three glioblastoma specimens (248_T, 259_T, 269_T; data from Ravi et al.[69]). EDNRB is consistently more highly expressed than EDNRA. **B**. Spatial transcriptomic expression maps (Ravi et al.[69]) for the same specimens, displaying EDNRA and EDNRB expression, endothelial and pericyte marker signatures, and the “Cluster 7” gene set from Wang et al. (2023) [68] with or without EDNRA. For each patient: (top left) H&E-stained section; (top middle) spatial gene expression maps (blue-to-red scale); (top right) spatial distribution of cell-type markers and Cluster 7 signature; (bottom left) spatial clustering of Visium spots; (bottom right) violin plots of expression across spatial clusters. EDNRA shows a more restricted pattern than EDNRB and is enriched in regions co-locating with vessel markers. These analyses indicate that tumor cells with the Cluster-7 signature and EDNRA expression share features with endothelial cells and pericytes.

## Acknowledgements

We acknowledge K. Ligon, O. Abigail (Dana-Farber Institute Harvard, US) and E. Huillard, N. Magne (ICM, Paris) for providing BT138, BT237 oligodendrogliomas cell lines. We acknowledge S. Weiss, A. Luchman (UofC, Calgary, Canada) for providing BT054 and BT088 oligodendrogliomas cell lines. We thank the MRI-MGX (supported from the France Génomique National Infrastructure (ANR-10-INBS-09). We acknowledge the Functional Proteomics Platform (S. Urbach, M. Seveno, S. Chaumont-Dubel, K. El Koulali) and the Montpellier Proteomics Platform (PPM, BioCampus Montpellier; ProFI UAR 2048, ANR-24-INBS-0015) for mass spectrometry analyses. We thank the Montpellier Vectorology Platform (MRI-PVM, C. Lemmers) for lentivirus production, and the MRI Imaging and Flow Cytometry Platform (A. Sarrazin, M.-P. Blanchard, V. Garcia) for technical support. We are grateful to the “Réseau d’Histologie Expérimentale de Montpellier” (RHEM) for tissue processing and histology, supported by SIRIC Montpellier Cancer (INCa_Inserm_DGOS_12553), the European Regional Development Fund, and the Occitanie region (FEDER-FSE 2014-2020). We acknowledge the MRI imaging facility (France-BioImaging, ANR-10-INBS-04) and the ARPEGE facility (L. Prezeau, D. Maurel, C. Vol, I. Brabet) for HTRF and ratiometric probe analyses. We are also really grateful to Cisbio-Revvity (E. Dupuis, E. Trinquet) for providing help, guidance and partnership with these.

## Author Contributions

**D Pineau** and **J-P. Hugnot** conceptualized the study and designed the methodology. **D. Pineau** performed all the experiments with the help of other coauthors. **L Garcia**, **H Arnold** and **A Hanžek** helped with cell culture, bulk RNA-seq, qPCR. **L Garcia** performed flow cytometry experiments, provided data analysis for endothelin receptors expressions and helped with all the figures design. **H Arnold** helped with western blotting and HUVEC cocultures experiments. **A Hanžek** performed qPCR for endothelin receptors expressions in hypoxia experiments. **S Hideg** and **K Aguilar-Cazarez** helped with primary cell cultures, western blotting and immunostaining experiments. **C Ripoll** helped with the patient sections and all immunostainings. **A Herbet, M Hautière** and **D Boquet** provided endothelin receptors antibodies notably RB49 monoclonal antibody, CHO-engineered cell lines and data analysis. **V Asei-Ceschino, L Gauthier, C Granotier-Beckers, F D Boussin** performed and analyzed migration assays on LGG275 cells. **S Urbach** and **M Seveno** helped with all mass spectrometry experiments from samples processing to data analysis and proteomic figures visualization. **C Jacques, L Brard, T Harnois, A Chatelier, V Coronas, B Constantin, J Chemin** provided Fura-2-AM calcium data and performed electrophysiological analyses of endothelin effects on SK2/SK3 channels (KCNN2/3). **P Rondard, L Prezeau, JP Pin** helped with EDNRB downstream signaling investigation using HTRF kits and ratiometric probes from experiments to data interpretation. **M Zheng, Guo-Hao Huang, Sheng-Qing Lv, L Zhang** provided help with patient sections, tumor microarray and data mining as for endothelin receptors explorations. **M Arrieta** provided spatial transcriptomics data mining. **H. Duffau** performed surgery on cases with diffuse low-grade gliomas. **L. Bauchet**. performed surgery on cases with glioblastomas. **V Rigau** provided the diagnostics and clinical information of the cases. **D Boquet, C Truillet, F Denat, H Duffau, J-P. Hugnot** secured funding support (Funding Acquisition). **J-P. Hugnot** and **D Pineau** wrote and corrected the manuscript. All authors reviewed and approved the final version of the manuscript.

