## Supplementary_Information for "Endothelin Signaling via EDNRB receptor Reduces Proliferation and Promotes Proneural-to-Mesenchymal Transition in Gliomas"

##### Public datasets (bulk and single-cell RNA-seq).

Expression of EDNRA and EDNRB were queried in bulk RNA-seq datasets (REMBRANDT, TCGA, CGGA) via the GlioVis portal [1], [2], [3], [4], [5] and in glioma single-cell RNA-seq datasets [6], [7], [8], [9] using the Gephart lab GBMseq browser and the Broad Institute Single Cell Portal [10].

##### Cell culture conditions.

Cultures were maintained at 37 °C in a humidified incubator with 5% $CO_2$ in DMEM/F12 (#21331046, Life Technologies) supplemented with L-glutamine (2 mM; #25030024, Thermo Fisher), N-2 supplement (1×; #17502048, Life Technologies), B-27 without vitamin A (1×; #12587010, Life Technologies), heparin (2 µg/mL; H3149-100KU, Sigma), EGF (10 ng/mL; AF-100-15-1MG, Peprotech), FGF2 (10 ng/mL; #100-18B-500UG, Peprotech), ciprofloxacin (10 µg/mL), gentamycin (10 µg/mL; #11520506, Thermo Fisher), and fungin (10 µg/mL; Invivogen). This medium is referred to as +GFs. The –GFs medium was identical except for the omission of heparin, EGF, and FGF2. Human glioblastoma stem cell lines were maintained as suspension cultures in vessels coated with poly(2-hydroxyethyl methacrylate) (poly-HEMA; 192066, Sigma-Aldrich). All other cell lines were cultured on vessels coated with poly-D-lysine (25 mg/mL, P7886, Sigma-Aldrich) and laminin (L2020, Sigma-Aldrich; hereafter referred to as PDL/Lam). Hypoxia experiments were performed using an incubator at 37°C in a humid atmosphere with 1% O2, (HeraCell Vios 160i). All cultures were routinely tested for the presence of mycoplasma. The identity of the cell cultures was verified by routine Short-Tandem Repeat (Eurofins). Cells were detached and collected after dissociation with Trypsin-EDTA 0.25% (#25200-056, Gibco), CaCl2 20 mM (10035-04-8, Fisher), DNAse I (#10104159001, Roche) and trypsin inhibitor (#17075029, Life Technologies). Cells were counted with a Z2 cell counter (Beckman Coulter). All experiments were carried out with cells below P15. For endothelin-response experiments (cell growth, proliferation, migration, apoptosis, transcriptomics, and proteomics), cells were cultured on Nunclon™ Delta tissue culture–treated plastic (Thermo Fisher Scientific) to minimize interference from laminin–integrin signaling. For CD44/OLIG1 flow cytometry following endothelin treatment (Fig. 4G) and for all signaling experiments using HTRF technology or fluorescent probes (Fig. 5), cells were cultured on poly-L-ornithine–coated surfaces (#P3655, 20 µg/mL; Sigma). CHO WT, CHO ETA (EDNRA-overexpressing), and CHO ETB (EDNRB-overexpressing) cell lines were generated and maintained as previously described [11].

##### RNA extraction, bulk RNA-seq and RT-qPCR.

Total RNA was extracted from cultured cells using the RNeasy kit (#74104, Qiagen) for RNA-seq, TRIzol (#15596018, Thermo Fisher), or the Arcturus PicoPure RNA kit (Thermo Fisher) for low cell numbers (e.g., FACS-purified cells). Patient tissues were homogenized in cold TRIzol using an Ultra-Turrax prior to extraction. RNA quantity was assessed with a NanoDrop spectrophotometer. For RT-qPCR, cDNA was synthesized from 200 ng RNA (low-grade cultures) or 500 ng RNA (glioblastoma/tissues) using random hexamers and reverse transcriptase, following the manufacturer's instructions. qPCR was performed with KAPA SYBR Fast (LC480 kit, #KK4610) on a LightCycler 480, and relative expression was calculated by the $2^{-\Delta\Delta Ct}$ method using ACTB as the reference gene. Primer sequences are listed in **Table S5**. For bulk RNA-seq, mRNA libraries were prepared and sequenced by BGI Genomics (Hong Kong, China or Poland) on the DNBSEQ™ platform (paired-end, 150 bp; ≥30 million reads/sample). Reads were quality-filtered with SOAPnuke, aligned to the reference genome with HISAT2, and quantified using RSEM to obtain counts, FPKM, and TPM values. Differential expression was analyzed with DESeq2 (FDR ≤ 0.05). Gene expression (TPM) data and volcano plots were generated using the BGI Dr. Tom platform.

##### Single-cell RNA sequencing analysis of LGG275 cell lines.

LGG275 (+GFs) grown in PolyHEMA-coated flasks were dissociated with trypsin-EDTA, filtered (40 µm), purified on a 15% Percoll gradient, and resuspended at 1,000 cells/µL in PBS/0.04% BSA. Viability was assessed with a CASY meter. Single-cell suspensions were processed on a Chromium Controller (10x Genomics) using the Chromium Single Cell 3' v3.1 kit, following the manufacturer's protocol. Libraries were sequenced on an Illumina NovaSeq 6000 (28 bp Read1, 10 bp I7, 10 bp I5, 90 bp Read2). Data were processed with Cell Ranger v6.1.1 (GRCh38), including a second --force-cells run to remove ambient RNA, aggregated with cellranger aggr, and visualized in Loupe Browser. Sequencing data are available at GEO (GSE263796). A detailed analysis of the single-cell RNA-seq results from LGG275 will be reported elsewhere (Garcia et al., in preparation).

##### Gene Set Enrichment Analysis (GSEA).

Transcriptomic profiles of glioma cell lines treated with endothelins were analyzed using GSEA (Dr. Tom platform (https://biosys.bgi.com) or GSEA software, [12], [13]) against the gene sets shown in the figures to identify enriched biological functions and pathways.

##### Protein extraction and Western blotting of patient tissues and cell cultures.

Cells and patient tissues were lysed in RIPA buffer supplemented with protease (cOmplete™ Ultra Tablets, #05892970001, Roche) and phosphatase inhibitors (PhosSTOP™, #04906845001, Roche). For patient tissues, homogenization was performed in 2 mL tubes containing 1.4 mm zirconium–silicate beads (#116913100, Lysing Matrix D, MP Biomedicals) using the manufacturer's protocol. Protein concentration was determined by BCA assay (#23225, #23227, Pierce™ BCA Protein Assay Kit, Thermo Fisher). For Western blotting, 10 µg of protein per sample was separated on 4–15% SDS–PAGE gels, transferred to Nitrocellulose or PVDF membranes, blocked, and incubated overnight at 4 °C with primary antibodies (see **Table S6**). HRP-conjugated secondary antibodies and ECL detection (Bio-Rad) were used; β-actin served as a loading control. Densitometry was performed using ImageLab v6.1 (Bio-Rad).

##### Immunofluorescence on cell cultures and patient tissue sections.

Immunofluorescence was performed on cells grown on glass coverslips and cultured in +GFs or –GFs medium for 5–7 days. For live staining, cells were incubated for 10 min at 4 °C in PBS containing 0.5% BSA and FcR blocking reagent followed by 15 min at room temperature with anti-EDNRB (RB49, 2 µg/mL). After three PBS washes, cells were fixed with 4% paraformaldehyde for 15 min at RT, permeabilized/blocked in PBS containing 5% donkey serum and 0.1% Triton X-100 for 30 min and labeled with Alexa Fluor secondary antibodies. For fixed-cell staining, fixation was performed first, followed by blocking/permeabilization and overnight incubation at 4 °C with anti-EDNRB and/or other primary antibodies (see **Table S6**). After PBS washes, Alexa Fluor secondary antibodies were applied for 1 h at RT. OCT-embedded patient samples were cut into 10 µm cryosections (SnapFrost®-frozen), fixed with 4% paraformaldehyde for 20 min at 4 °C, permeabilized/blocked (0.3% Triton X-100, 10% donkey serum), and incubated overnight at 4 °C with primary antibodies. Secondary antibodies were applied for 1 h at RT. Isotype controls and omission of primary antibodies were used as negative controls. All samples were counterstained with Hoechst 33342, mounted in Fluoromount (F4680, Sigma), and imaged using a Zeiss Apotome microscope (40–63×).

##### Flow cytometry.

For **Fig. S4**, LGG275 cells ($5 \times 10^5$) were cultured on PDL/laminin in +GFs or –GFs medium for 5–7 days, then labeled either live or after 4% paraformaldehyde fixation with anti-EDNRB (sheep) or RB49 (mouse) antibodies (2 µg/mL), followed by Alexa Fluor 647–conjugated secondary antibodies (1:2000). Fixed cells were permeabilized in PBS containing 1% BSA, 0.1% Triton X-100, and 0.5 mM EDTA. For CD44/OLIG1 labeling after endothelin treatment (Fig. 4G), LGG275 cells were plated on poly-L-ornithine–coated vessels in +GFs or –GFs medium and treated once daily for 5 days with either endothelin dilution buffer (control) or ET-1 (10 nM). Live cells were first labeled with PE-conjugated anti-CD44 (#130-110-394, Miltenyi), then fixed, permeabilized and stained with anti-OLIG1 (#AF2417, R&D). Nuclei were counterstained with Hoechst 33342. All samples were analyzed on a MACSQuant cytometer (405, 488, and 635 nm lasers; V1 450/50BP for Hoechst, R1 692/75 for Alexa Fluor 647) using isotype controls (rabbit IgG #3900, Cell Signaling; goat IgG #AB-108-C, Bio-Techne; sheep IgG #5-001-A, R&D; mouse IgG #X093101-2, Dako) and fluorescence minus one controls.

##### Cell growth assay.

Glioblastoma stem cells (20,000/well) or low-grade diffuse glioma cells (30,000/well) were seeded in 48-well plates with +GFs medium (400 µL/well). After adhesion, cells were treated every 2 days with endothelin dilution buffer (control) or recombinant ET-1 or ET-3 (4 nM, Sigma). The EDNRB agonist IRL1620 (10 nM, SCP0135, Sigma) was applied following the same schedule. Dose–response assays (0.01–100 ng/mL; 4 pM–40 nM) were also performed for ET-1 and ET-3 (**Fig. 3C**). After 15–20 days, cells were dissociated with Trypsin-EDTA and counted using a Z2 counter (n = 12 technical replicates per condition).

##### EdU incorporation and cell death assays.

LGG275 cells ($3\times10^4$/well) were seeded in 48-well plates and treated every 2 days for 5–6 cycles with control buffer, ET-1 (10 nM), ET-3 (10 nM), or IRL1620 (10 nM). Proliferation was assessed using the Click-It EdU kit (#BCK-EdUFC647, BaseClick) with 10 µM EdU (50 h), followed by fixation, permeabilization, Alexa Fluor 647–azide staining, and Hoechst 33342 counterstain. Cell death was evaluated using YO-PRO-1™/propidium iodide or Annexin V–AF647 labeling in appropriate buffers. Samples were analyzed on a MACSQuant cytometer (405, 488, 635 nm lasers) with isotype and fluorescence minus one controls.

##### Cell migration assay.

LGG275 cells ($3\times10^4$/well) were plated in laminin-coated 24-well plates (0.55 µg/cm², LN521, Biolamina). After 24 h, medium was replaced with endothelin-containing medium (10 nM), and time-lapse imaging was performed using a Nikon A1R confocal microscope under controlled conditions (37 °C, 5% $CO_2$, 19% $O_2$, 95% RH) as described previously [14]. Images were acquired in mosaic mode (8×8, 20× objective, zoom 2) every 10 min for 4 or 24 h. Migration velocity and plot-to-origin graphs were generated from 100 and 50 tracked cells per condition, respectively, using MTrackJ (ImageJ) and DiPer software [15].

##### Proteomic Analyses

Proteins from LGG275 cells ($1.5\times10^6$) (+GFs) and treated once daily for 5 days with vehicle (PBS + 0.1% BSA) or ET-1 (4 nM) were extracted in RIPA with protease/phosphatase inhibitors 8 h after the last treatment. Proteins were digested with trypsin (S-Trap, ProtiFi) and labelled with a TMT10 kit (Thermo Fisher). Pooled samples were fractionated (8 fractions) by basic RP chromatography (Pierce™ High pH Reversed-Phase Peptide Fractionation Kit) and analyzed by nanoLC–MS/MS (RSLC U3000 coupled to an Exploris480 with FAIMS). A gradient consisting of 0–31% B for 120 min, 31–50% B for 5 min (A = 0.1% formic acid, 2% acetonitrile in water; B = 0.1% formic acid in 80 %acetonitrile) at 300 nL/min was used to elute peptides from the capillary (0.075 mm×500 mm) reverse-phase column (Pepmap®, Thermo Fisher Scientific). Data were processed with MaxQuant v2.0.3.0 using the

RefProteome_HUMAN-cano_2023_02_UP000005640 and contaminant databases. Fixed modification: carbamidomethyl (C); variable: oxidation (M), acetyl (protein N-term) and FDR ≤ 1% at protein/peptide level. Significance thresholds were determined using Perseus (version 1.6.15.0) based on the Significance B approach, with a p-value cutoff set at 0.01 for protein ratios (Et/Ctrl). Only proteins found to be significantly regulated in at least 3 out of 5 replicates were retained for further analysis.

The mass spectrometry proteomics data have been deposited to the ProteomeXchange Consortium via the PRIDE [16], partner repository with the dataset identifier PXD065902 and 10.6019/PXD065902.

##### $IP_1$ Production and ERK1/2 Phosphorylation Assays (HTRF)

$IP_1$ accumulation and ERK1/2 phosphorylation were quantified using the HTRF IP-One Gq kit (#62IPAPE, Cisbio) and the HTRF Phospho-ERK1/2 kit (#64ERKPEG, Cisbio), respectively, following the manufacturer's protocols [17], [18]. For $IP_1$ assays, LGG275 cells were seeded in Poly-L-ornithine–coated plates in +GFs or –GFs medium and cultured for 7 days. Cells were then treated for 5 min with vehicle (PBS + 0.1% BSA), ET-1 or ET-3 (40 nM), the EDNRB agonist IRL1620 (0-1000 nM), or the EDNRB antagonist BQ-788 (0-100 000 nM) in the presence of $EC_{80}$ ET-1 (4 nM). Cells were lysed in the kit lysis buffer and incubated with $d_2$-coupled $IP_1$ antibodies and terbium cryptate–coupled anti-$IP_1$. For $pERK_1/_2$ assays, cells were lysed in saturation buffer containing protease and phosphatase inhibitors, then incubated with Europium cryptate–coupled and $d_2$-coupled anti-$pERK_1/_2$ antibodies. TR-FRET signals (665 nm/620 nm ratio) were recorded using a PHERAstar plate reader. $IP_1$ concentrations were calculated from the kit's standard curve, and $pERK_1/_2$ values were expressed relative to control conditions.

##### Intracellular Calcium Measurements

For population-based measurements, LGG275 cells ($1 \times 10^5$) were seeded in Poly-L-ornithine–coated 96-well plates (+GFs/–GFs, 7 days) and loaded with Cal520-AM (1.25 µM, 1 h, 37 °C) in PBS containing 0.5 mM $CaCl_2$ and 0.5 mM $MgCl_2$. Fluorescence responses to ET-1, ET-3, IRL1620 agonist, BQ-788 antagonist were recorded at 37 °C using an FDSS µCell plate reader (Hamamatsu Photonics; Ex 493 nm/Em 515 nm). ΔF (max–min) values were integrated and normalized to control with *FDSS Analyzer V3*. For single-cell ratiometric imaging, LGG275 cells ($5 \times 10^4$) on Poly-D-lysine/laminin–coated glass coverslips were loaded with Fura-2-AM (6 µM, 30 min, 37 °C, 5% $CO_2$) and imaged in external buffer (130 mM NaCl, 5.4 mM KCl, 0.8 mM $MgCl_2$, 10 mM HEPES, 5.6 mM glucose, 1.8 mM $CaCl_2$, pH 7.4). Excitation at 340 and 380 nm with emission at 505 nm was acquired every 2 s at 37 °C using a Lambda 421–equipped Olympus IX73 microscope with Zyla sCMOS camera and *Metafluor* software. The 340/380 nm fluorescence ratio was calculated over time from selected region of interest and normalized to baseline.

#### Measurement of potassium fluxes

Potassium channel activity was assessed in LGG275 cells using the FluxOR probe (#F20015, ThermoFisher) according to the manufacturer's protocol. After a 1-min baseline recording, cells were stimulated with ET-1 (40 nM), ET-3 (40 nM), the EDNRB agonist IRL1620 (1 µM), or ET-1/IRL1620 with apamin (1 µM), and fluorescence was recorded for 5 min on an FDSSµcell (Hamamatsu Photonics) at 37 °C. Data were analyzed with FDSS Analyzer V3 to quantify potassium fluxes.

#### Patch clamp

LGG275 cells ($5.10^3$) were seeded in PDL/Lam coated 35mm-diameter Petri dish 7 days before experimentation. For potassium channel identification, Petri dishes were also coated with sDLL4 to further promote a non-proliferative state as previously . The measurements were carried out at room temperature. Patch electrodes (3–5 M) were pulled from borosilicate glass capillaries (GC-150T, Harvard Aparatus) using a vertical micropipette puller (Narishige, Tokyo, Japan). Voltage-clamp experiments in whole-cell configuration were performed using an Axopatch 200B amplifier with a CV 203BU headstage (Molecular Devices, CA). Voltage command pulses were generated by a personal computer equipped with an analog-digital converter (Digidata 1550; Molecular Devices) using pCLAMP software v11.0 (Molecular Devices). Currents were filtered at 5 kHz and digitized at 10 kHz. The digitized currents were stored on a computer for later offline analysis. Intrapipette solution was composed of KCl 145 mM, MgCl2 1 mM, Mg-ATP 1 mM, HEPES 10 mM, CaCl2 8.7 mM, EGTA 10 mM, pH 7.2. The final pCa was 6 ([free $Ca^{2+}$] = 1 µM) in experiments designed to detect the presence of SK channels by direct stimulation with intracellular solution. In experiments using an extracellular ligand (endothelin) to induce a physiological intracellular $Ca^{2+}$ response a PCa 9 ([free $Ca^{2+}$]= 1 nM) was used with the following intrapipette solution: KCl 145 mM, MgCl2 1 mM, Mg-ATP 1 mM, HEPES 10 mM, EGTA 0.5 mM, pH 7.2. The extracellular solution consisted of NaCl 140 mM, KCl 4 mM, $MgCl_2$ 1 mM, $CaCl_2$ 2 mM, glucose 11.1 mM, HEPES 10 mM, pH 7.4. Macroscopic currents were measured in response to a ramp protocol, from -100mV to +60mV for 4s from a resting potential of 0mV. The interval between each ramp was 5 seconds. To monitor the effect of Apamin and ET1 on current amplitude, cells were held in the standard bath solution and perfused with this control solution to measure the control current. Then, the same solution containing either Apamin, ET1 or ET1+Apamin was applied to the cell using a microperfusion system. Data were analysed using a combination of pCLAMP software v11.0 (Molecular Devices), Microsoft Excel, and GraphPad Prism 8.0.

#### Effect of Cytokines, YAP and Notch Signaling on EDNRA and EDNRB Expression

LGG275 cells were treated for five consecutive days with selected cytokines and pathway modulators. Treatments included ET-1 (10 ng/mL, 4 nM), OSM (10 ng/mL), LIF (10 ng/mL), BMP2 (10 ng/mL), BMP4 (10 ng/mL), BMP6 (10 ng/mL), BMP7 (10 ng/mL), IFNγ (10 ng/mL), and TDI-011536 (5 µM) targeting the YAP1 pathway. Medium containing the respective compounds was renewed once daily. Cells were collected 8 h after the final treatment for RNA or protein extraction (See **Tables S1, S4-5**). For virus-mediated pathway modulation, LGG275 or Gb7 cells were transduced with lentiviral or adenoviral vectors to overexpress constitutively active STAT3 (STAT3C; MOI = 50; gift from Linzhao Cheng, Addgene plasmid #24983;[19]), constitutively active YAP (YAP1-S5A; MOI = 10; in-house generated adenovirus;[20]), or activated Notch (NICD;[21]), together with the corresponding control vectors. After a 15-day expansion period, transduced cells were sorted by FACS to isolate positive populations. Total RNA was extracted (Arcturus Picopure RNA kit) and processed for quantitative PCR analysis. Protein lysates were prepared in RIPA buffer supplemented with protease and phosphatase inhibitors for Western blotting.

#### Glioma Cell–HUVEC Co-culture, RNA Extraction and Bulk RNA Sequencing

LGG275 (GFP-labelled) and Gb7 (YFP-labelled) glioma cells, as well as HUVEC endothelial cells (mCherry-labelled), were generated by lentiviral transduction followed by FACS sorting. One day before co-culture, $3.4 \times 10^5$ HUVECs (Human Umbilical Vein Endothelial Cells; Promocell) were seeded on gelatin-coated plates in complete endothelial growth medium (#C-22111, Promocell) supplemented with recombinant human EGF (5 ng/mL), basic FGF (10 ng/mL), IGF Long R3 (20 ng/mL), $VEGF_{165}$ (0.5 ng/mL), ascorbic acid (1 µg/mL), heparin (22.5 µg/mL), hydrocortisone (0.2 µg/mL), and fetal calf serum (2% v/v). Glioma cells ($3.0 \times 10^5$) were then added and co-cultured for 48 h. Following co-culture, $GFP^+$ or $YFP^+$ glioma cells and $mCherry^+$ HUVECs were isolated by FACS. Total RNA was immediately extracted using the Arcturus Picopure RNA kit (Thermo Fisher) for bulk RNA sequencing.

#### Spatial transcriptomics data analysis in glioblastoma from Ravi et al

The 10X Visium spatial transcriptomic dataset from Ravi et al. was downloaded from Datadryad (https://datadryad.org/stash/dataset/doi:10.5061/dryad.h70rxwdmj) and analyzed using the Seurat R package. From the full cohort, samples UKF248T, UKF 259T and UKF269 were selected for our analyses since they showed the most interesting spatial expression levels and patterns of all samples.

Spatial clustering was performed using the BayesSpace algorithm (https://www.nature.com/articles/s41587-021-00935-2). The endothelial cell signature was defined via the following genes: PECAM1, CDH5 and KDR, whereas for the pericyte signature the markers PDGFRB, MYH11, NOTCH3, HIGD1B, RGS5 and ACTA2 were considered. Signatures were calculated using the AddModuleScore function from the Seurat package. Spatial expression visualizations were generated via the SpatialFeaturePlot function and violin plots were created via the VlnPlot function.

#### Statistical analysis

For all experiments, the statistical methods, number of replicates, and number of independent experiments are specified in the corresponding figure legends. Data analyses and graph generation were performed with GraphPad Prism software, version 10.0.0 (153) (GraphPad Software, Boston, MA, USA). Significance thresholds were defined as follows: ns (not significant), $p \geq 0.05$; * (significant), $0.01 \leq p < 0.05$; ** (very significant), $0.001 \leq p < 0.01$; *** (extremely significant), $0.0001 \leq p < 0.001$; **** (very extremely significant), $p < 0.0001$.

#### Data availability

All raw and processed scRNA-seq data generated in this study have been deposited in the Gene Expression Omnibus (GEO) under accession number GSE263796. All raw and processed RNA-seq data generated in this study have been deposited in the Gene Expression Omnibus (GEO) under accession number such as GSE298185 (for LGG275 treated with compounds), GSE298282 (for glioma cell lines upon ET-1), GSE298356 and GSE298357 (for LGG275 in coculture with endothelial cells) and GSE298358 (for Gb7 in coculture with endothelial cells).

### Supplementary Tables

**Table S1**: Lists of all products and reagents

| Name | Reference | Final concentration | Fabricant |
|---|---|---|---|
| DMEM/F12 | #21331046 | / | Life Technologies |
| L-glutamine | #25030024 | 2 mM | Thermo Fisher |
| B-27 without vitamin A (50X) | #12587010 | 1X | Life Technologies |
| N-2 (100X) supplement | #17502048 | 1X | Life Technologies |
| rhEGF | AF-100-15-1MG | 10 ng/mL | Peprotech, ThermoFisher |
| rhFGF2/bFGF | 100-18B-500UG | 10 ng/mL | Peprotech, ThermoFisher |
| heparin | H3149 | 2 µg/mL | Life Technologies |
| gentamicin | #11520506 | 10 µg/mL | Sigma-Merck |
| ciprofloxacin | #17850 | 10 µg/mL | Fisher Scientific |
| fungin | #ant-fn-1 | 10 µg/mL | Sigma-Merck |
| Poly-HEMA<br>2-hydroxyethyl methacrylate | #192066 | 10 mg/mL | Invivogen |
| Hypoxia incubator | / | / | HeraCell Vios 160i |
| Trypsin-EDTA | #25200-056 | 0.25% | Life Technologies |
| DNAse I | #10104159001 | 10 mg/µl | Roche |
| $CaCl_2$ | #10035-04-8 | 20 mM | Fisher Scientific |
| Trypsin inhibitor | #17075029 | 50 mg/mL | Life Technologies |
| Z2 cell counter | #Z2 9914591-D | / | Beckman Coulter |

| | | | |
|---|---|---|---|
| RNEasy kit | #74104 | / | Qiagen |
| TRIzol, | #15596018 | / | Thermo Fisher |
| KAPA SYBR Fast LC480 PCR kit | #KK4610 | / | Promega |
| Ultra-Turrax Disperser IKA | (RNA_extracts) | / | ThermoFisher |
| FastPrep-24 | (Protein_extracts) | / | MP_Biomedical |
| cOmplete™ Ultra Tablets | #05892970001 | / | Roche |
| PhosSTOP™, | #04906845001 | / | Roche |
| Pierce™ BCA Protein Assay Kit | #23225, #23227 | | Thermo Fisher |
| Laemmli loading buffer | / | 1 X | Sigma Aldrich |
| Amersham Protran nitrocellulose membranes (0.20 µm-pore size) | / | / | GE Healthcare |
| Mini-PROTEAN TGX Stain Free Gels 4-15%, | #4568086 | / | Bio-Rad |
| Ladder Kaleidoscope | #1610375 | / | Bio-Rad |
| EveryBlot BlockingBuffer | #12010020 | / | Bio-Rad |
| anti-rabbit IgG-HRP-linked | / | 1/2000 | GE Healthcare |
| Secondary HRP-linked antibodies | / | 1/2000 | Jackson Laboratory |
| Clarity Western ECL Substrate | #170-5060 | / | Bio-Rad |
| ChemiDoc MP Imaging System | / | / | Bio-Rad |
| Immobilon ECL | #WBULS0100 | / | Millipore-Sigma |
| GeneGnome XRQ Imaging System | / | / | Syngene |
| Poly-L-Ornithine | #P3655 | 1X | Millipore-Sigma |
| Nunclon Delta surface | #140685 | / | ThermoFisher |
| anti-Annexin V-AF647 antibody | #640912, | / | BioLegend |
| YO-PRO-1™ | #V13243 | / | ThermoFisher |
| Click-It EdU kit | #BCK-EdUFC647 | / | BaseClick |
| FcR blocking reagent | #130-059-901 | / | Miltenyi Biotec |

| | | | |
|---|---|---|---|
| TMT10 kit | #90110 | / | Thermo Fisher |
| nanoLC–MS/MS (RSLC U3000 + Exploris480 with FAIMS) | / | / | Thermo Fisher |
| HTRF IP-One Gq kit | #62IPAPE, | / | Cisbio, Revvity |
| HTRF Phospho-ERK1/2 kit | #64ERKPEG | / | Cisbio, Revvity |
| PheraStar | / | / | BMG, Labtech |
| Cal520-AM | #Cal-520® AM | 1,25 µM | AATBioquest |
| Fura-2AM | #F1201 | 6 µM | ThermoFisher |
| FluxORTM | #F20015 | / | ThermoFisher |
| FURA2-AM Lambda 421–equipped Olympus IX73 microscope with Zyla sCMOS camera | / | / | Olympus, Oxford instruments |
| FDSS µcell | / | / | Hamamatsu Photonics |

| **Name** | **Reference** | **Stock concentration** | **Dilution** | **Final concentration** | **Fabricant** |
|---|---|---|---|---|---|
| ET-1 | E7764-10UG | 4µM (10µg/mL) | PBS+BSA 0,1% | Defined in the study | Sigma |
| ET-3 | E9137-10UG | 4µM (10µg/mL) | PBS+BSA 0,1% | Defined in the study | Sigma |
| IRL-1620 | SCP0135-500UG | 100 µM | DMSO | Defined in the study | Sigma |
| BQ-788 | HY-15894A | 10 mM | DMSO | Defined in the study | MedChemExpress |
| hOSM | 300-10 | 10 µg/mL | PBS+BSA 0,1% | 10 ng/mL | Peprotech |
| hLIF | 300-05 | 10 µg/mL | PBS+BSA 0,1% | 10 ng/mL | Peprotech |
| hCNTF | 450-13 | 10 µg/mL | PBS+BSA 0,1% | 10 ng/mL | Peprotech |
| hBMP2 | 120-02C | 10 µg/mL | PBS+BSA 0,1% | 10 ng/mL | Peprotech |
| hBMP4 | 120-05 | 10 µg/mL | PBS+BSA 0,1% | 10 ng/mL | Peprotech |
| hBMP6 | 120-06 | 10 µg/mL | PBS+BSA 0,1% | 10 ng/mL | Peprotech |
| hBMP7 | 120-03P | 10 µg/mL | PBS+BSA 0,1% | 10 ng/mL | Peprotech |
| hIFNβ | 300-02BC | 10 µg/mL | PBS+BSA 0,1% | 10 ng/mL | Peprotech |
| hIFNγ | 300-02 | 10 µg/mL | PBS+BSA 0,1% | 10 ng/mL | Peprotech |
| TDI-011536 | HY-150042 | 5 mM | DMSO | 5 µM | MedChemExpress |

**Table S2** : List of patient specimens

| Patient samples name | Patient gender | Patient Age | Diagnosis (subype, grade) | IDH1 status | Extracted from biological samples | Use for which experiments (EDNRA/EDNRB) | IF/IHC |
|---|---|---|---|---|---|---|---|
| Gb39 | F | 50 | GB | IDH WT | Protein | Western Blot | |
| Gb40 | F | 81 | GB | IDH WT | RNA, protein | RT-qPCR, Western Blot | |
| Gb43 | M | 52 | GB | IDH WT | RNA, protein | RT-qPCR, Western Blot | |
| LGG321 | M | 28 | A2 | IDH1R132H | RNA, protein | RT-qPCR, Western Blot | |
| LGG330 | M | 37 | A2 | IDH1R132H | RNA, protein | RT-qPCR, Western Blot | |
| LGG250 | F | 40 | A2/3 | IDH1R132H | RNA, protein | RT-qPCR, Western Blot | |
| LGG348 | F | 25 | A3 | IDH1R132H | RNA, protein | RT-qPCR, Western Blot, Cryosections | EDNRB/APOE |
| LGG309 | F | 61 | A3 | IDH1R132H | RNA | RT-qPCR | |
| LGG309 (2) | F | 61 | A3 | IDH1R132H | RNA | RT-qPCR | |
| LGG356 | M | 39 | A3 | IDH1R132H | RNA, protein | RT-qPCR, Western Blot | |
| LGG322 | M | 52 | O2/3 | IDH1R132H | RNA | RT-qPCR | |
| LGG346 | M | 45 | O2/3 | IDH1R132H | RNA, protein | RT-qPCR, Western Blot, Cryosections | EDNRB/IDH1R132H, Ctrl |
| LGG358 | M | 43 | O2 ou O3 | IDH1R132H | RNA, protein | RT-qPCR | |
| LGG351 | M | 60 | O3 | IDH1R132H | RNA, protein | RT-qPCR, Western Blot | |
| LGG357 | M | 29 | O2 (/O3) | IDH1R132H | RNA, protein | RT-qPCR, Western Blot, Cryosections | EDNRB/IDH1R132H |
| LGG357bis | M | 29 | O2 (/O3) | IDH1R132H | RNA | RT-qPCR | EDNRB/APOE ; EDNRB/OLIG2 |
| LGG359 | F | 62 | O3 | IDH1R132H | RNA, protein | RT-qPCR, Western Blot | |
| LGG361 | M | 40 | O3 | IDH1R132H | Protein | Western Blot, Cryosections | EDNRB/IDH1R132H, Ctrl |
| LGG362 | M | 25 | O3 | IDH1R132H | Protein | Western Blot | |
| LGG318 | M | 26 | A2 | IDH1R132H | Protein | Western Blot | |
| LGG309 | F | 61 | A3 | IDH1R132H | Protein | Western Blot | |
| LGG355 | M | 44 | A4 | IDH1R132H | Protein | Western Blot | |
| LGG244 | M | 40 | A2 | IDH1R132H | Sections | Cryosections | EDNRB/OLIG2 |
| Gb34-A | M | 60 | GB | IDH WT | Protein | Western Blot | |
| LGG 180 (180-1) | F | 35 | O2 | IDH1R132H | Protein | Western Blot | |
| LGG184 (184-O) | F | 38 | A2 | IDH1R132H | Protein | Western Blot | |
| LGG187 (187-1) | M | 26 | A2 | IDH1R132H | Protein | Western Blot | |
| LGG182 (182) | M | 25 | A2/3 | IDH1R132H | Protein | Western Blot | |
| LGG190 (190-2) | M | 42 | A2/3 | IDH1R132H | Protein | Western Blot | |
| LGG189 (189-2) | F | 27 | A3 | IDH1R132H | Protein | Western Blot | |
| LGG93 | N.A | N.A | O3 | IDH1R132H | Protein | Western Blot | |
| LGG185 (185) | M | 39 | A4 | IDH1R132H | Protein | Western Blot | |
| LGG56 (56) | M | 33 | A4 | IDH1R132H | Protein | Western Blot | |
| LGG85 (85) | M | 38 | A4 | IDH1R132H | Protein | Western Blot | |
| LGG316 | M | 59 | O2 | IDH1R132H | Protein | Western Blot | |
| Meningioma | N.A | N.A | / | / | Protein | Western Blot | |
| Human Brain | N.A | N.A | / | / | Protein | Western Blot | |

(A= Astrocytomas, O=Oligodendrogliomas, GB = Glioblastomas)

**Table S3**: Glioma tissue microarray [22], [23].

| | Case no | Gender | Age | Diagnosis | WHO grade | Core 1 | Core 2 |
|---|---|---|---|---|---|---|---|
| **1** | 3 | F | 37 | Anaplastic astrocytoma | 3 | 0 | 0 |
| **2** | 4 | F | 57 | Diffuse astrocytoma | 2 | 1 | 0 |
| **3** | 6 | M | 40 | Anaplastic oligodendroglioma | 3 | | |
| **4** | 7 | F | 65 | Oligodendroglioma | 2 | | |
| **5** | 26 | M | 57 | Control WM & GM | | 1 | 1 |
| **6** | 27 | M | 34 | Anaplastic oligoastrocytoma | 3 | 1 | 0 |
| **7** | 33 | M | 54 | Glioblastoma | 4 | 1 | 2 |
| **8** | 34 | F | 67 | Anaplastic astrocytoma | 3 | 1 | 1 |
| **9** | 35 | M | 83 | Diffuse astrocytoma | 2 | 1 | 0 |
| **10** | 37 | F | 70 | Anaplastic oligodendroglioma | 3 | | |
| **11** | 38 | M | 29 | Oligodendroglioma | 2 | | |
| **12** | 55 | M | 28 | Control WM & GM | | 1 | 0 |
| **13** | 56 | M | 49 | Anaplastic oligoastrocytoma | 3 | 1 | 2 |
| **14** | 60 | M | 65 | Glioblastoma | 4 | 0 | 1 |
| **15** | 61 | F | 31 | Anaplastic astrocytoma | 3 | 0 | 0 |
| **16** | 62 | M | 27 | Diffuse astrocytoma | 2 | 0 | 0 |
| **17** | 64 | F | 60 | Anaplastic oligodendroglioma | 3 | | |
| **18** | 76 | M | 38 | Control WM & GM | | 0 | 0 |
| **19** | 77 | F | 61 | Anaplastic oligoastrocytoma | 3 | 1 | 0 |
| **20** | 80 | F | 29 | Anaplastic astrocytoma | 3 | 0 | 0 |
| **21** | 81 | M | 32 | Diffuse astrocytoma | 2 | 0 | 0 |
| **22** | 83 | F | 42 | Anaplastic oligodendroglioma | 3 | | |
| **23** | 84 | F | 65 | Oligodendroglioma | 2 | | |
| **24** | 92 | F | 24 | Anaplastic oligoastrocytoma | 3 | 1 | 0 |
| **25** | 93 | F | 65 | Glioblastoma | 4 | 2 | 2 |
| **26** | 94 | F | 46 | Anaplastic astrocytoma | 3 | 2 | 2 |
| **27** | 95 | M | 48 | Diffuse astrocytoma | 2 | 0 | 0 |
| **28** | 102 | M | 34 | Anaplastic oligoastrocytoma | 3 | 0 | 0 |
| **29** | 103 | F | 57 | Glioblastoma | 4 | 1 | 2 |
| **30** | 104 | M | 41 | Anaplastic astrocytoma | 3 | 0 | 1 |
| **31** | 105 | F | 42 | Diffuse astrocytoma | 2 | 0 | 0 |
| **32** | 106 | F | 34 | Anaplastic oligodendroglioma | 3 | | |
| **33** | 112 | F | 63 | Glioblastoma | 4 | 2 | 2 |
| **34** | 113 | F | 37 | Anaplastic astrocytoma | 3 | 1 | |
| **35** | 114 | M | 26 | Diffuse astrocytoma | 2 | 1 | 1 |
| **36** | 115 | M | 42 | Anaplastic oligodendroglioma | 3 | | |
| **37** | 116 | M | 42 | Glioblastoma | 4 | 1 | 1 |
| **38** | 117 | F | 64 | Glioblastoma | 4 | 2 | 2 |
| **39** | 120 | M | 55 | Glioblastoma | 4 | 0 | 2 |

Annotation: 0 = no vascular staining; 1 = minority of vessels stained; 2 = majority of vessels stained

#### Table S4: Characteristics of the glioma cell lines

| Cell lines | Patient (Sex-Age) | Tumor information | Grade (WHO 2016) | IDH1 status | Miscellaneous information | Alterations | Publication status |
|---|---|---|---|---|---|---|---|
| LGG275 | F40 | Recurrent primary Tumor (reoperation of 2014) | Grade II / III astrocytoma | IDH1R132H | Astrocytoma grade II according to histology astrocytoma high grade according to molecular annotation | ATRX lost Presence in 3 copies of EGFR, MYC, Braf, c-Met, loss of one copy of CDKN2A | Augustus et al, 2021 |
| LGG336 | F40 | Recurrent primary tumor (reoperation of 2012) | Grade II / III astrocytoma | IDH1R132H | Astrocytoma grade II according to histology | c-Met and Braf 3 copies, loss of 9q | *Unpublished (Garcia et al, in preparation)* |
| LGG85 | M38 | Recurrent primary tumor (reoperation of 2012) | Grade IV astrocytoma | IDH1R132H | TMZ | c-Met neg | Leventoux et al, 2020 |
| LGG349 | F57 | Recurrent primary Tumor (reoperation of 2018) | Grade IV astrocytoma | IDH1R132H initially but mutation loss in culture | multiple cycles of PCV, radiotherapy + TMZ | ATRX maintained MGMT methylation, unmutated TERT | *Unpublished (Garcia et al, in preparation)* |
| BT138 | M50 | Recurrent primary Tumor (reoperation) | Grade III Oligodendroglioma (NOS) | IDH1R132H initially but mutation loss in culture | Tumor in the left frontal lobe | | Koivunen et al, 2012 |
| BT237 | F43 | Recurrent primary Tumor (reoperation) | Grade III Oligodendroglioma | IDH1R132H | Tumor in the left frontal lobe | | Koivunen et al, 2012 |
| BT054 | F49 | Recurrent primary Tumor | Grade III Oligodendroglioma | IDH1R132H | TMZ and radiotherapy Tumor in the right frontal lobe | MGMT methylation | Kelly et al, 2010; Luchman et al, 2012; Mazor et al, 2015; Yuan et al, 2018 |
| BT088 | M50 | Recurrent primary Tumor (reoperation) | Grade III Oligodendroglioma | IDH1R132H initially but mutation loss in culture | Tumor in the right frontal lobe Several cycles of PCV, radiotherapy+TMZ | | Kelly et al, 2010; Luchman et al, 2012; Mazor et al, 2015; Yuan et al, 2018 |
| Gb4 | M53 | Primary Tumor | Grade IV Glioblastoma | IDH1 WT | Mesenchymal | | Guichet et al, 2013 Guichet et al, 2015 |
| Gb5 | M64 | Primary Tumor | Grade IV Glioblastoma | IDH1 WT | GB with oligodendroglial component | | Guichet et al, 2013 Guichet et al, 2016 |
| Gb7 | M52 | Primary Tumor | Grade IV Glioblastoma | IDH1 WT | Proneural | | Guichet et al, 2013; Guichet et al, 2015 |
| Gb21 | F53 | Primary Tumor | Grade IV Glioblastoma | IDH1 WT | Giant Cell GB | | Guichet et al, 2016; Guelfi et al, 2021 |

**Table S5 :** List of RT-qPCR primers

| **Gene** | **Forward Primer** | **Reverse Primer** |
|---|---|---|
| ACTB | GGACTTCGAGCAAGAGATGG | AGCACTGTGTTGGCGTACAG |
| **EDNRA** | TCAAGATGGAAACCCTTTGC | ATTGAGCCATTGCTGGGTAG |
| **EDNRB** | ATGACGCCACCCACTAAGAC | GAACACAAGGCAGGACACAA |
| CD44 | CAATAGCACCTTGCCCACAAT | AATCACCACGTGCCCTTCTATG |
| CTGF | CAGCATGGACGTTCGTCTG | AACCACGGTTTGGTCCTTGG |
| OLIG1 | CGCAGAGAGTTTTCGCTCTT | GCGGTTGGTTTTCGTTTTTA |
| OLIG2 | GACAAGCTAGGAGGCAGTGG | CGGCTCTGTCATTTGCTTCT |
| MKI67 | CCCCCACCAGAACTAACAGA | ACTTTGATGCCCTCATCACC |
| STAT3 | CAGCAGCTTGACACACGGTA | AAACACCAAAGTGGCATGTGA |
| NOTCH1 | TCCACCAGTTTGAATGGTCA | CGCAGAGGGTTGTATTGGTT |
| KCNN3 | CTGCCGCCAAAATAAACATT | GCCTGGCACAAGCTTTCTAC |

**Table S6 :** List of antibodies

| Name | Species | Reference | Manufacturer | Dilution |
|---|---|---|---|---|
| EDNRA | Rabbit | ab178454 | Abcam | 1:1000 |
| EDNRB | Sheep | AF4496-SP | R&D Systems | 1:500 |
| RB49 | Mouse | CEA | from Dr Boquet & Dr Herbet | 1:500 |
| OLIG1 | Goat | AF2417 | R&D Systems | 1:300 |
| OLIG2 | Mouse | MABN50 | Sigma | 1:500 |
| ASCL1 | Rabbit | ab211327 | Abcam | 1:100 |
| GFAP | Rabbit | Z0334 | Dako | 1:5000 |
| AQP4 | Rabbit | 16473-1-AP | Proteintech | 1:500 |
| APOE | Rabbit | EP1374Y | Abcam | 1:1000 |
| pSMAD1/5 | Rabbit | 9516 | Cell Signalling | 1:800 |
| IDH1R132H | Mouse | DIA−H09 | Dianova | 1:100 |
| CD44-PE | Rabbit | 130-110-394 | Miltenyi Biotec | 1:100 |
| CTGF | Rabbit | Ab209780 | Abcam | 1:500 |
| pSTAT3 | Rabbit | 9145S | Cell Signalling | 1:1000 |
| STAT3 | Rabbit | A19566 | ABClonal | 1:1000 |
| pERK1/2 | Rabbit | 9910 | Cell Signalling | 1:1000 |
| ERK1/2 | Rabbit | 9102 | Cell Signalling | 1:1000 |
| KCNN3/SK3 | Rabbit | H-45 | Santa Cruz Biotechnology | 1:100 |
| KCNN2/SK2 | Rabbit | APC-028 | Alomone Labs | 1:200 |
| α-actin | Rabbit | A2066 | Sigma-Aldrich | 1:5000 |
| β-actin | Mouse | 8H10D10 | Novus | 1:2500 |
